# Inflammasome Activation and IL-1β Release in Alveolar Macrophages Infected with *Pseudomonas aeruginosa* is Reduced in Hypoxia

**DOI:** 10.64898/2026.09.14.750667

**Authors:** Alexander W. Rapp, Aurora L. Golden, Arianna D. Reuven, Jay Goddard, Emily Ann McClure, Aimee M. Wilson, Chloe A. Young, Caitlin E. Murphy, Taalia-Lindsay Morgan, Andrew J. Olive, Joshua J. Obar, Deborah A. Hogan, Benjamin D. Ross, James B. Bliska

**Author notes:** Corresponding author: James B. Bliska Geisel School of Medicine at Dartmouth Department of Microbiology & Immunology 524A Remsen, 66 College Street Hanover, NH 03755.

## Abstract

*Pseudomonas aeruginosa* is an important agent of acute or chronic airway infections. A complex set of genetic and environmental factors determine outcomes of *P. aeruginosa* airway infections.

*P. aeruginosa* undergoes genetic adaptation to finetune expression of a type III secretion system (T3SS) and other virulence factors in response to selective pressures during infection. Genetic and environmental factors predispose for *P. aeruginosa* infections in patients such as those with cystic fibrosis (CF). CF is a genetic disorder resulting in hypoxia and thick mucus that fosters *P. aeruginosa* airway infection. Resident alveolar macrophages (AMs) coordinate immune responses to *P. aeruginosa* by producing proinflammatory cytokines such as IL-1β, which can promote infection resistance. AMs infected with *P. aeruginosa* laboratory strains detect the T3SS and activate the NLRC4 inflammasome, resulting in IL-1β release. Studies of inflammasome responses in AMs to a *P. aeruginosa* CF isolate of have not been reported. Here, we characterized the CF isolate DH1137 and found it has a downregulated but functional T3SS, is adapted to grow in hypoxia, and induces significant production of the inflammasome cytokines IL-1α and IL-1β during acute lung infection in a CF mouse model. Inflammasome gene expression was primed in AMs by LPS, and upon DH1137 infection these cells released IL-1β. Infection of AMs with DH1137 or laboratory strain PAO1 in hypoxia resulted in significantly dampened inflammasome activation compared to normoxia. These data describe a new mechanism that *P. aeruginosa* may exploit to evade immune detection by tissue-resident lung macrophages in the context of CF.

**Importance:** Airway infection of people with CF by *P. aeruginosa* results from genetic and environmental changes that impact host and pathogen. Mutations in the cystic fibrosis transmembrane conductance regulator gene (*CFTR*) cause anatomical alterations in airways, including areas of hypoxia, that compromise anti-bacterial defense. *P. aeruginosa* genetically adapts by downregulating a T3SS important for establishment and turning up phenotypes that promote chronic infection such as hypoxic growth. AMs are resident airway phagocytes that can normally detect *P. aeruginosa and* undergo inflammasome activation to release IL-1 cytokines, however these cells are known to be dysregulated in CF, and their interactions with clinical isolates are understudied. Here show that *P. aeruginosa* elicits IL-1 cytokines during acute lung infection in a CF mouse model but triggers low inflammasome activation in hypoxic AMs ex vivo, suggesting the bacterium exploits low oxygen to evade detection by resident lung macrophages in CF.

## Introduction

*Pseudomonas aeruginosa* is a Gram-negative opportunistic bacterial pathogen that is an important agent of acute or chronic airway infections (1). The outcome of a *P. aeruginosa* airway infection is determined by a complex set of genetic and environmental factors. *P. aeruginosa* has numerous virulence factors including flagellar motility, a type III secretion system (T3SS), and an alginate exopolysaccharide that gives it a mucoid phenotype and is a major component of biofilms (1). *P. aeruginosa* can undergo genetic adaptation to fine tune the expression of these and other virulence factors in response to selective pressures encountered during airway infection (1). Genetic and environmental factors predispose many patient groups for *P. aeruginosa* airway infections that are associated with high morbidity and mortality, including individuals with ventilator-associated pneumonia, chronic obstructive pulmonary disease, or cystic fibrosis (CF) (1, 2). Epithelial, resident alveolar macrophages (AMs) and recruited phagocytes (e.g. neutrophils) and other cells coordinate immune responses to *P. aeruginosa* in airways (2). Airway innate immune responses to *P. aeruginosa* result in proinflammatory cytokine production, which can be important for infection resistance, but excessive inflammation can be damaging and promote pathogen persistence (2–4). Metabolites and other products of metabolism produced by airway phagocytes and other organs such as liver (e.g. ketone bodies) or gut (e.g. short chain fatty acids) can accumulate in the lungs, impacting infection and promoting disease tolerance to *P. aeruginosa* (5–7).

CF is an autosomal recessive disorder caused by mutations in the Cystic Fibrosis Transmembrane Conductance Regulator (*CFTR*) gene, which encodes a crucial chloride and bicarbonate ion channel. These mutations correlate with profound immune dysregulation and a high susceptibility to intermittent (childhood) and chronic (adulthood) *P. aeruginosa* airway infections (8, 9). CF airways feature areas of acidosis, hypoxia, and thick mucus that fosters bacterial infections, particularly chronic *P. aeruginosa* biofilms. As a primary driver of airway infections, *P. aeruginosa* contributes significantly to morbidity in people with CF (pwCF) (10, 11), with roughly 50–70% of adults developing long-term, persistent infections (12).

Furthermore, aberrant inflammation is a critical mediator of CF pathogenesis and closely parallels progressive lung function decline (13). While the advent of highly effective CFTR modulator therapies (HEMT) has dramatically improved the lifespan and overall well-being of pwCF (14–17), *P. aeruginosa* continues to persist and adapt in the airway post treatment (18). Recent clinical data also indicate that persistent airway inflammation from *P. aeruginosa* infection continues to drive lung damage even post-treatment (19).

PwCF exhibit markedly elevated levels of the proinflammatory cytokine IL-1β in airway secretions (20–22). Once released, IL-1β acts directly on airway epithelial cells to upregulate mucin genes, compounding airway obstruction (20). Elevated IL-1β is observed in bronchoalveolar lavage fluid (BALF) even in the absence of detectable infection (21).

Experimental airway infections using mice carrying mutations in the *Cftr* gene have confirmed inflammatory responses including IL-1β production in response to *P. aeruginosa* (23–26). Studies have reported that transcripts or protein for IL-1β are elevated during *P. aeruginosa* infection in the CF murine model as compared to control mice (27).

This overabundant IL-1β release highlights the central role of host inflammasome activation in CF pathophysiology (20). Mechanistically, inflammasome protein complexes are vital components of innate immune surveillance that detect the presence of pathogen molecules in the host cell cytosol. Upon detecting cytosolic pathogen-associated molecular patterns (PAMPs), inflammasome receptors assemble, serving as a platform for caspase-1-dependent Gasdermin D (GSDMD) cleavage and pore formation to facilitate the secretion of mature IL-1β and IL-18, which signal for the recruitment and activation of additional antibacterial immune cells to clear the infection (28).

*P. aeruginosa* has a T3SS that delivers effectors into host cells and is central to both virulence (29, 30) and the inflammasome response to infection (2–4, 31, 32). The T3SS is comprised of a basal body in the bacterial cell envelope and a needle tipped with the PcrV protein. Upon contact with a host cell, proteins secreted through the needle form a channel in the plasma membrane, to allow translocation of effectors. Three T3SS effectors, ExoT, ExoU and ExoS, are important virulence factors (30, 33). ExoS is produced by 58-72% of clinical *P. aeruginosa* isolates and *exoS^+^* strains cause most CF infections (34, 35). The translocation of effectors by the *P. aeruginosa* T3SS is accompanied by the delivery of needle, rod and flagellin subunits into the host cell cytosol which are recognized by a NAIP co-receptor for the NLRC4 inflammasome (31, 32). Activation of the NLRC4 inflammasome in murine cells is triggered when an NAIP recognizes and binds to the T3SS needle (NAIP1), rod (NAIP2) or flagellin (NAIP5/NAIP6) proteins (31, 32). Activation of the NLCR4 inflammasome in macrophages infected with *P. aeruginosa* results in release of IL-1β and cell death by pyroptosis (36–38). The effectors ExoU or ExoS can reduce inflammasome activation in phagocytes infected with *P. aeruginosa* in certain contexts (4, 37, 39–43). For example, ExoS inhibits NLRC4 inflammasome activation in neutrophils (39, 41, 42) but not macrophages (36–38) infected with *P. aeruginosa*. *P. aeruginosa* isolates from established airway infections in pwCF have typically undergone genetic adaptation toward phenotypes important for chronic persistence such as decreased T3SS function and increased biofilm capacity (10, 44–46), and these strains typically fail to activate inflammasomes in infected macrophages (40). However, a recent report showed that an acute infection of human CF or non-CF monocyte-derived macrophages (MDMs) with the *P. aeruginosa* mucoid clinical isolate DH1137 resulted in release of IL-1β, IL-18 and IL-1α (47), suggesting that some chronic strains retain sufficient T3SS function to be detected by the NLRC4 inflammasome.

The phenotypic and functional diversity of macrophages is heavily influenced by their microenvironment, driven largely by their capacity to adapt to low-oxygen niches. Within the CF airway, a steep oxygen gradient exists within the thickened mucus, creating a hypoxic environment shared by both host macrophages and *P. aeruginosa* (48–50). The persistence of *P*.

*aeruginosa* in the CF lung is directly aided by its ability to survive within this hypoxic mucus as a biofilm. At the molecular level, the hypoxia-inducible factor (HIF) family of transcription factors critically regulates how macrophages adapt to oxygen deprivation (51–53). Indeed, studies using murine bone marrow-derived macrophages (BMDMs) or human macrophages exposed to hypoxia report enhanced inflammasome activation across multiple disease contexts (54–56). Despite these insights into oxygen-deprived cellular mechanisms, how bacterial pathogens influence macrophage inflammasome activation under hypoxic conditions remains underexplored. While recent work by Okano et al. reported that hypoxia enhances NLRP3 inflammasome activation in BMDMs infected with the Gram-negative anaerobic bacterial pathogen *Porphyromonas gingivalis* (57), whether inflammasome activation by tissue-resident lung macrophages in response to *P. aeruginosa* is similarly altered within the hypoxic CF environment remains unknown.

Recent research has identified tissue-resident alveolar macrophages (AMs) as central players in regulating pulmonary inflammation. Under homeostatic conditions, AMs serve as vital immune sentinels in the lungs, maintaining tissue integrity while surveying for foreign pathogens.

However, human primary CF AMs exhibit a paradoxically dysregulated phenotype: they display a heightened inflammatory response that correlates with impaired bacterial clearance (58, 59).

Single-cell transcriptomic analyses by Schupp et al. showed that pre-HEMT the majority of macrophages in the human CF sputum are derived from recruited monocytes from the periphery (60). More recently, analysis of BALF cells from pwCF on HETM showed that AMs are the most abundant macrophages, and that these cells display transcriptomic changes as compared to healthy controls (61). Among these alterations, single-cell RNA sequencing identified distinct AM clusters from BALF of pwCF on HEMT with signatures of heightened inflammasome- related inflammation compared to healthy controls (61).

AMs possess a distinct developmental lineage, seeding the lungs from the fetal liver during embryonic development (62–64). Orchestrated by the local cytokines GM-CSF and TGF-β, AM differentiation results in a phenotype that is uniquely hypoinflammatory against many pathogenic stimuli compared to BMDMs (64–68). Consequently, standard BMDM cultures fail to recapitulate these key functional nuances of the lung. To overcome this limitation, Olive et al. recently developed an *ex vivo* murine model termed fetal liver-derived alveolar-like macrophages (FLAMs) (69). By culturing fetal liver cells in the presence of GM-CSF and TGF-β, this method yields a stable, self-replicating population that closely mirrors primary AM functions and phenotypes (69). The FLAM model has been utilized to study Gram-positive *Mycobacterium abscessus* infections (70). We sought to leverage this system to evaluate tissue-resident macrophage inflammasome responses to *P. aeruginosa*.

Laboratory strains of *P. aeruginosa* have been shown to activate the NLRC4 inflammasome in murine AMs in vitro (38) and in vivo (71, 72). However, studies of inflammasome responses in AMs to a clinical CF isolate of *P. aeruginosa* have not been reported. Here, we characterized the mucoid clinical isolate DH1137 and found that it has a functional T3SS, is adapted to grow in hypoxia, and induces significant production of the inflammasome cytokines IL-1α and IL-1β during lung infection in a CF mouse model. Inflammasome gene expression was primed in FLAMs by LPS stimulation, and upon DH1137 these cells underwent GSDMD cleavage and IL- 1β release. By directly comparing FLAMs against both BMDMs and primary alveolar macrophages isolated from mouse lungs (AlvMacs), we demonstrated that FLAMs behave similarly to primary AlvMacs yet differ substantially from BMDMs regarding inflammasome activation by DH1137. Intriguingly, we found that infection of FLAMs with DH1137 in hypoxia resulted in significantly dampened inflammasome activation in comparison to normoxia. These data describe a novel use of the FLAM model to study infection by a clinical Gram-negative bacterial isolate, providing critical new insights into how *P. aeruginosa* evades immune detection by tissue-resident lung macrophages in the context of CF.

## Results

### DH1137 has a Functional T3SS, is Motile and Adapted to Hypoxia

DH1137 has been used as a CF clinical isolate in several published studies (47, 73–75). Recently, Aridgides et al. showed that CF and non-CF MDMs infected with DH1137 release IL-1 cytokines (47), suggesting that this strain has a functional T3SS that is detected by the NLRC4 inflammasome.

To determine if DH1137 has a functional T3SS overnight cultures of this isolate and the control *P. aeruginosa* laboratory strain PAO1 were subcultured and grown to an OD_600_=0.5, without or with EGTA to induce T3SS expression and secretion. Western blot analysis of intracellular and TCA-precipitated extracellular fractions revealed that DH1137 produced and secreted ExoS, although at substantially lower levels than PAO1 (**Figure S1A**). We sequenced DH1137 and a related *lasR* mutant isolate DH1136 (Materials and Methods). Comparison of DH1137 with PAO1 revealed several codon changes in genes known to regulate T3SS expression (e.g. *exsD,* **Table S1**). In addition to these codon changes that could reduce T3SS expression, we noted that DH1137 has a T120A codon change in *mucA* that may be associated with its high mucoid and low T3SS phenotypes (76) (**Table S1**). *P. aeruginosa* clinical isolates are known to adapt to hypoxia in the CF airway (77). We investigated if T3SS production and secretion are altered in *P. aeruginosa* in response to a switch from normoxia (21% O2) to hypoxia (1% O2). In hypoxia, intracellular ExoS levels in DH1137 decreased compared to normoxia, while secreted amounts remained unchanged (**Figure S1A**). In contrast, production and secretion of ExoS increased in PAO1 in hypoxia (**Figure S1A**). The normoxia/hypoxia comparison for DH1137 was repeated and we additionally Western blotted for the T3SS tip protein PcrV. Reduced production of ExoS in hypoxia was confirmed, but this effect did not extend to PcrV (**Figure S1B**). To examine how T3SS induction and hypoxia impacted bacterial replication, we monitored growth via OD600 readings. Addition of EGTA decreased growth of DH1137 and PAO1 overall (**Figure S2**).

Surprisingly, DH1137 grew more efficiently in hypoxia than in normoxia, while the opposite was seen for PAO1 (**Figure S2**). Codon changes in the DH1137 genome that could be associated with adaptation to hypoxia have not been defined but include mutations in the *anr* or *dnr* genes. Overall, these results indicate that DH1137 is adapted to hypoxic growth and confirms that reduced ExoS production under this condition is not an artifact of decreased bacterial replication.

In addition to T3SS loss of function mutations, selective pressures within the CF airway frequently drive loss of bacterial flagellin, resulting in non-motile phenotypes (78). Early studies tracking host-adapted strains found that over 40% of clinical isolates from people with CF (pwCF) completely lack flagella or display severe swimming defects (79, 80). To compare DH1137 and PAO1 motility phenotypes, overnight cultures were subcultured without or with EGTA, spotted onto swimming plates, and incubated under normoxic or hypoxic conditions. A PAO1 *ΔfliC* mutant was used as a negative control. After 14 hours in normoxia, DH1137 exhibited swimming motility that was substantially greater than PAO1 (**Figure S3A**).

Furthermore, motility assays revealed that oxygen depletion drastically reduced DH1137 swimming (**Figure S3A**) or swarming after 18 hours (**Figure S3B**). Together, these findings demonstrate that DH1137 is positive for T3SS, adapted to grow in hypoxia, and motile, the extent to which is strongly regulated by oxygen availability.

### DH1137 Lung Infection Promotes Inflammation Dominated by IL-1 Cytokines in a CF Mouse Model

To our knowledge DH1137 has not been studied in a mouse infection model. To define the virulence characteristics of DH1137 during lung infection, we utilized CF mice which have the F508del mutation in Cftr (81). F508del heterozygous (F508del-Het) and homozygous (F508del) mice were born germfree and vertically engrafted with a synthetic gut microbiota (SGM) community (82) (see Materials and Methods). Prior to infection the F508del mice had decreased abundance of Bacteroides in their SGM compared to F508del-Het (**Figure S4**), reflecting the dysbiosis of pwCF (83–85). F508del-Het and F508del mice were inoculated with DH1137 intratracheally. Control mice were inoculated with PBS (naïve). Mouse weights were monitored over five days. F508del-Het and F508del mice showed similar and significant reductions in weights, reaching ∼15% loss compared to starting (**Figure 1A**). At the two-day post infection timepoint, a separate cohort of mice was euthanized for both CFU and cytokine analysis of their lungs. The day 2 timepoint was chosen as the day to perform CFU analysis, as this matches with the day at which the most significant weight loss was seen. CFUs of recovered bacteria from the lung homogenates trended higher in F508del compared to F508del-Het but were not significantly different and showed a roughly 10-fold reduction compared to the starting inoculum (**Figure 1B**). Past findings utilizing CF mice for acute lung infection with planktonic *P. aeruginosa* noted rapid declines in bacterial burden (86, 87). Lung homogenates from the infected and naive mice were additionally utilized for cytokine analysis via a Luminex^®^ assay. Among 32-tested, the inflammasome-related cytokines IL-1α and IL-1β had the highest significance differences comparing naïve to all infected (**Figure S5**). IL-1β levels trended higher in F508del compared to F508del-Het but the differences were not significant (**Figure 1C**). G-CSF, which is produced by macrophages and signals to neutrophils, was significantly higher in F508del compared to F508del-Het (**Figure 1D**). These data show that DH1137 induces lung inflammation in a CF mouse infection model and that the acute inflammatory response is largely driven by Il-1family cytokines. The robust *in vivo* production of IL-1β following DH1137 infection strongly points toward the activation of host inflammasome complexes in lung macrophages.

**Fig. 1.**
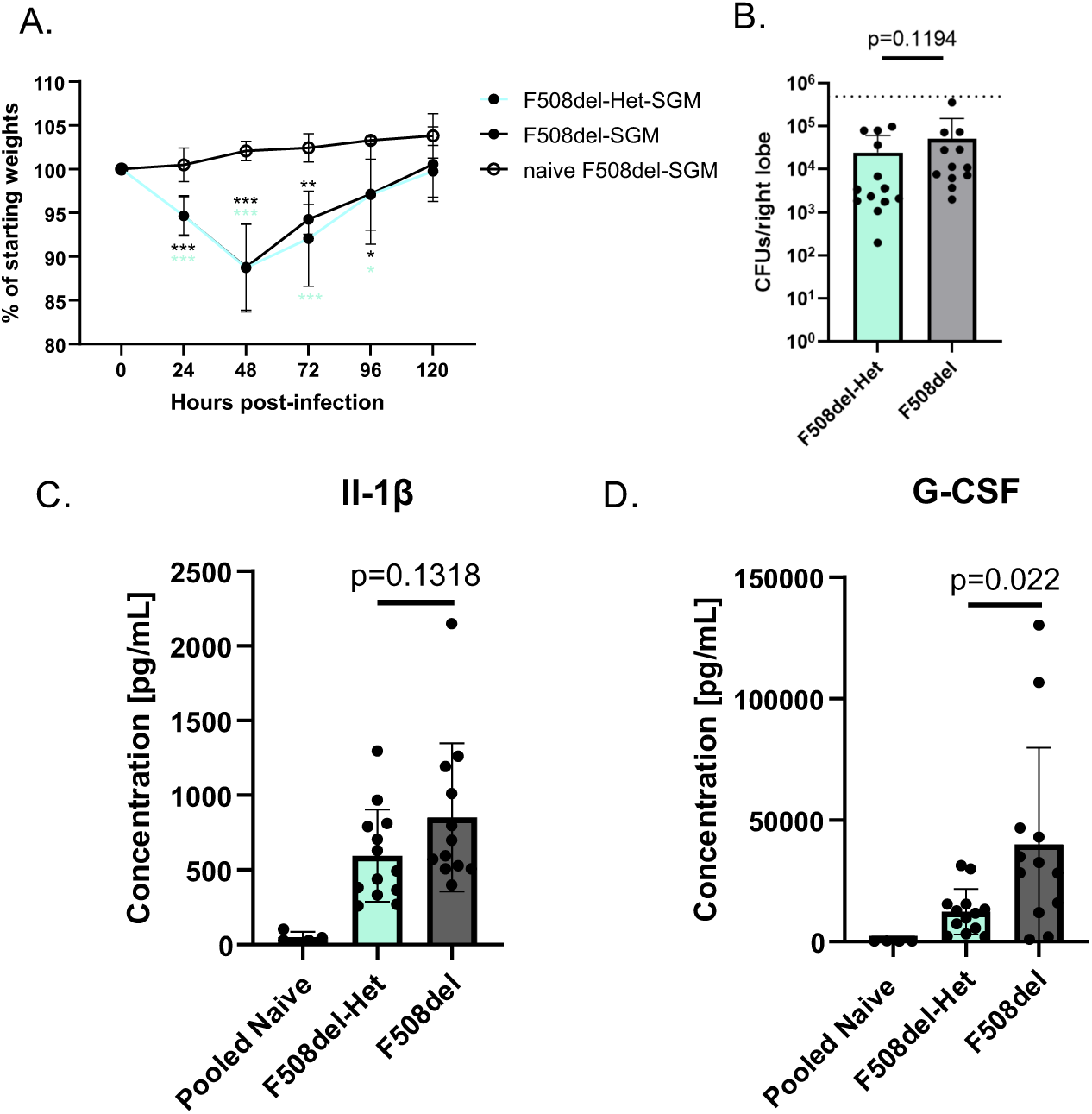
**DH1137 infection outcomes in F508del-Het-SGM or F508del-SGM mice.** 508del-Het or F508del mice (both female and male) were inoculated intratracheally with x10^5^ CFU of DH1137 or PBS alone (naïve). (A) Percent of starting body weight over ime. Error bars are means and standard deviations of pooled data from 3 independent xperiments, n=2 for naïve, n=18 for F508del-Het and n=9 for F508del. Differences in eights between time 0 and later points were tested for significance by paired t-test **p<0.001; **p<0.01; *p<0.05. (B) Log10 CFU per right lung lobes at 48 hrs post- nfection pooled from 3 independent experiments, n=13 for F508del-Het and n=12 for 508del. Error bars are means and standard deviations. Dotted line shows the noculation dose. Significance determined by Mann-Whitney test. (C,D) Cytokine levels etermined by 32-Plex Luminex^®^ assay using the same lung homogenates analyzed in anel B with additional naïve controls (F508del, F508del-Het or WT) combined, n=4. onparametric Mann-Whitney tests were used to determine p-values between the 508del-Het and F508del groups.

### Fetal Liver-Derived Alveolar Macrophages Undergo Inflammasome Activation Upon Infection with DH1137

A recent study reported that AMs are the most abundant cell type in BALF from pwCF with mild to moderate lung disease and currently on HEMT (61). In addition, gene expression analysis indicated that inflammasome-related genes including Il1b were upregulated in these AMs (61).

FLAMs have been shown to release IL-1β following priming with IFNγ and subsequent *Mycobacterium abscessus* infection, indicating inflammasome pathway activation in these cells (70). To understand if inflammasome responses to Gram negative bacteria can be primed in these cells, we performed bulk RNA sequencing (RNA-seq) on FLAMs stimulated with *E. coli*- derived LPS for 18 hours. Differential expression analysis identified 2,046 upregulated and 2,075 downregulated genes following LPS exposure (**Figure S6A**). Notably, many genes involved in inflammasome pathways were significantly upregulated (e.g. Il1b, Nlrp3, Casp1, Naip2). Gene ontology (GO) analysis of the top enriched biological processes revealed pathways having to do with cellular responses to bacteria and response to molecules of bacterial origin (**Figure S6B**). **Figure S6C** shows a heat map of relevant gene expression standardized by Z-score. Together, these data demonstrate that FLAMs can robustly respond to LPS priming by upregulating key inflammasome-related genes.

We next infected LPS-primed FLAMs with DH1137 at MOI 10 for 1.5 hrs and measured outcomes. Prior to infection DH1137 was subcultured without or with EGTA, the later representing T3SS-inducing conditions. Western blotting of cell lysates showed that priming increased production of pro-IL-1β and Hif1α in FLAMs (**Figure 2A**). Hif1α is known to be produced in response to LPS stimulation in addition to hypoxia in macrophages (88, 89), and increased expression of Hif1α was detected by RNA-seq in primed FLAMs (**Figure S6AC**). Infection with DH1137 promoted GSDMD cleavage, which was greater with T3SS-induced bacteria (**Figure 2A**). Corresponding with GSDMD cleavage in response to DH1137 infection, there were significant increases in mature IL-1β secretion as quantified by ELISA (**Figure 2B**). Interestingly, cytotoxicity as measured by LDH release was significantly increased upon DH1137 infection but did not differ between T3SS inducing conditions (**Figure 2C**). Viability of DH1137 as measured by CFU assay indicated that T3SS inducing conditions resulted in significantly increased growth of the bacteria (**Figure 2D**), likely due to ExoS antiphagocytic activity (90). Consistent with CFU results, phase contrast video microscopy of infected FLAMs revealed that T3SS inducing conditions resulted in greater numbers of extracellular motile DH1137 (**Video S1AB**). Notably, “swarms” of motile T3SS-induced DH1137 could be seen in the vicinity of individual FLAMs (**Video S1AB**). Together, these data suggest that the T3SS in DH1137 is detected by the NLRC4 inflammasome in LPS-primed FLAMs, resulting in GSDMD pore formation and release of mature IL-1β.

**Fig. 2.**
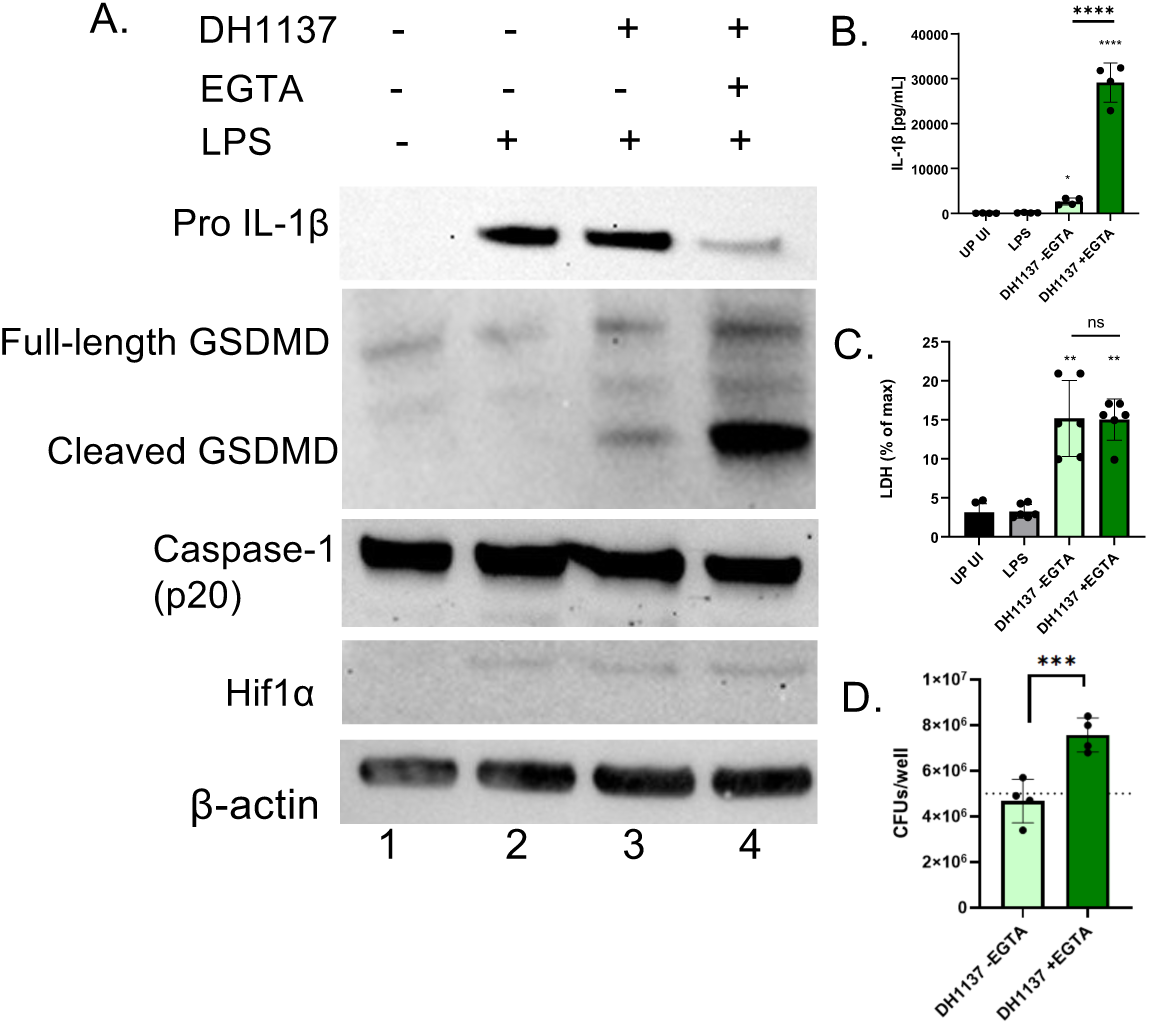
DH1137 infection outcomes in FLAMs. FLAMs were left unprimed (UP) or primed with LPS for 18 hrs and then left uninfected (UI) or infected for 1.5 hrs at a MOI=10 with DH1137 following subculture in the absence or presence of EGTA . (A) Western blotting of FLAM lysates was performed to probe for the levels of the indicated proteins. Results are representative of 3 Independent experiments. (B) IL-1β levels in supernatants were quantified via ELISA. (C) FLAM pyroptosis was quantified as percentage of max LDH release in supernatants. (D) Viable bacteria in the combined supernatant and cellular fraction was quantified by CFU assay, with input of 5x106. In B-D, data is pooled from 4 or more independent experiments where each dot represents a single well, and bars show the mean ±SD. Statistical significance in (B,C) was determined using one-way ANOVA and Tukey post test comparing to LPS alone or between conditions as shown by brackets. In (D) statistical significance determined by student’s *t*-test (****p<0.0001, ***p<0.001, **p<0.01, ns not significant).

### Comparison of Infection Responses to *P. aeruginosa* between FLAMs and BMDMs

To compare the responses of FLAMs infected with DH1137 to another type of macrophage we used bone marrow-derived macrophages (BMDMs). BMDMs were isolated from C57BL/6J mouse femurs and tibias and differentiated with M-CSF according to standard protocols (91). FLAMs and BMDMs were seeded at identical densities for direct comparison. Following overnight LPS priming, Western blot analysis revealed that BMDMs produced less pro-IL-1β, more HIF1α and equivalent NLRP3 as compared to FLAMs (**Figure 3A**). The Hif1α result aligns with prior studies showing that this protein is maintained at higher basal levels in BMDMs than in tissue-resident AMs to regulate steady-state metabolic function (92). Upon DH1137 infection BMDMs displayed greater GSDMD cleavage, lower IL-1β release, and higher cytotoxicity compared to FLAMs (**Figure 3A,B,C**). BMDMs were also less efficient than FLAMs at controlling DH1137 replication, especially under T3SS-inducing conditions (**Figure 3D, Video S1CD**).

**Fig. 3.**
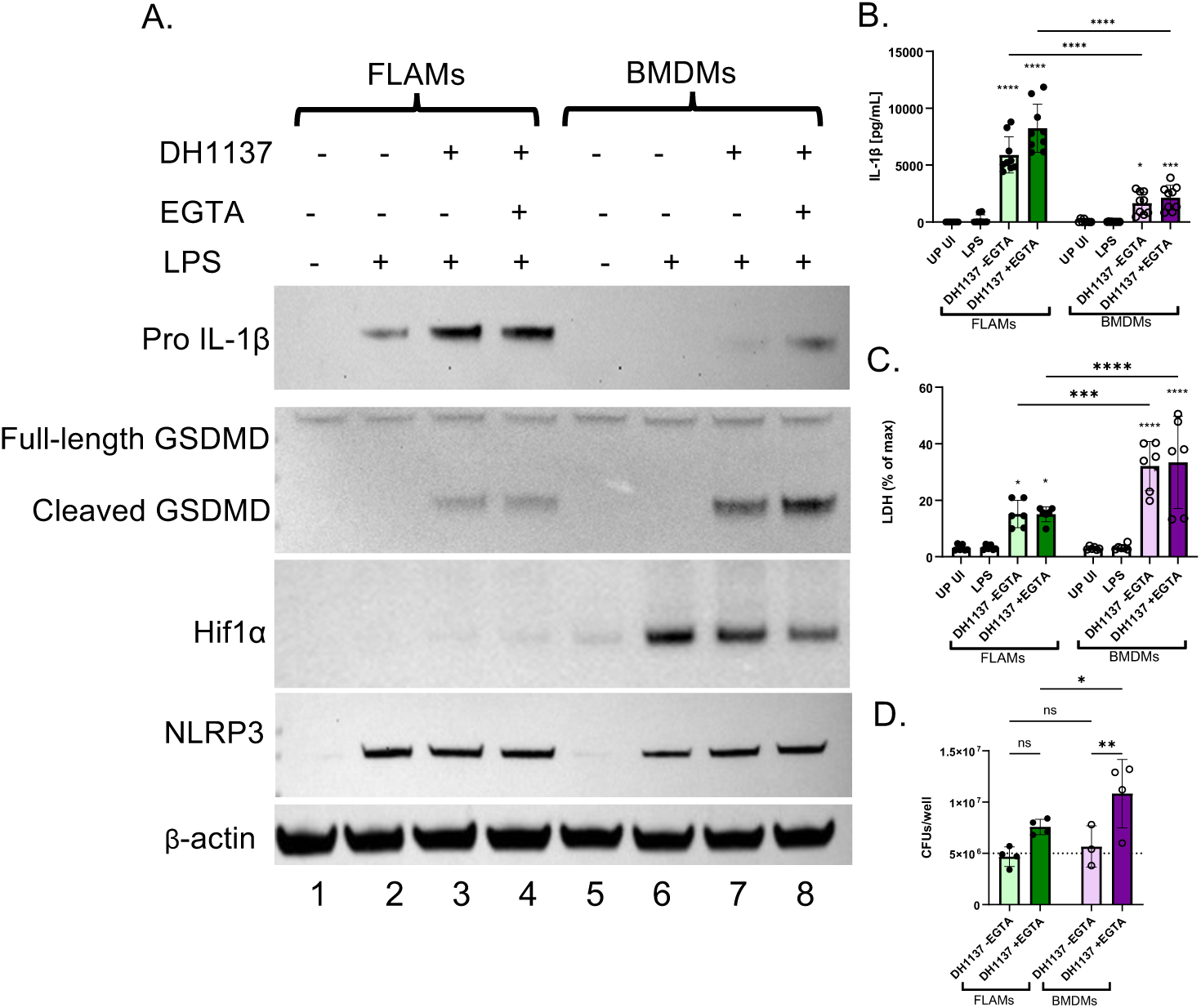
DH1137 infection outcomes in FLAMs or BMDMs. FLAMs or BMDMs were left UP or primed with LPS for 18 hrs and then left UI or infected a MOI=10 for 1.5 hours with DH1137 following subculture in the absence or presence of EGTA. (A) Western blotting of FLAM or BMDM lysates was performed to probe for the levels of the indicated proteins. Results are representative of 3 Independent experiments. (B) IL-1β levels in supernatants were quantified via ELISA. (C) FLAM and BMDM pyroptosis was quantified as a percent of maximum LDH release. (D) Viable bacteria in the combined supernatant and cellular fraction was quantified by CFU assay, with input of 5x10^6^. In B-D data is pooled from 3 or more threeindependent experiments where each dot represents a single well, and bars show the mean ±SD. Statistical significance in (B,C) was determined using two-way ANOVA and Sidaks multiple comparisons post test comparing to LPS alone or between conditions as shown by brackets. In (D) statistical significance was determined using a student’s *t*-test. (****p<0.0001, ***p<0.001, **p<0.01, *p<0.05, ns not significant).

Inflammasome response differences between FLAMs and BMDMs infected with PAO1 were also investigated. While the decrease in IL-1β release in BMDMs vs FLAMs was less pronounced following infection with PAO1, it remained statistically significant (**Figure S7A**). BMDMs also displayed compromised infection control, yielding higher PAO1 CFUs relative to FLAMs (**Figure S7B, Video S2**). Overall, these results indicate that FLAMs and BMDMs exhibit differences in their priming and infection responses to *P. aeruginosa* infection, with the former having higher pro-IL-1β production, mature IL-1β release and bactericidal activity and lower cytotoxicity.

### Comparison of Infection Responses to *P. aeruginosa* between FLAMs and AlvMacs

We next compared FLAMs against primary alveolar macrophages (AlvMacs) isolated from the BALF of C57BL/6J mice. Following *ex vivo* expansion, AlvMacs were seeded at identical densities to FLAMs, primed with LPS, and infected with T3SS-induced DH1137 as above except at MOI 1. The lower MOI was used to better match the lower seeding density. Western blotting showed similar priming responses for pro-IL-1β, Hif1α and NLRP3 between FLAMs and AlvMacs (**Figure 4A**). Upon DH1137 infection similar GSDMD cleavage was noted, although ELISA analyses revealed higher amounts of mature IL-1β were released from FLAMs as compared to AlvMacs (**Figure 4A,B**). FLAMs were also better equipped at controlling DH1137 replication (**Figure 4C, Video S3**). Results of using PAO1 in the FLAM/AlvMac comparison showed similar cytotoxicity and bactericidal activity but higher IL-1β release from the former (**Figure S7C,D,E**). Altogether, these data reveal that FLAMs align more closely with primary AlvMacs than BMDMs in terms of priming and inflammasome responses to *P. aeuruginosa*.

**Fig. 4.**
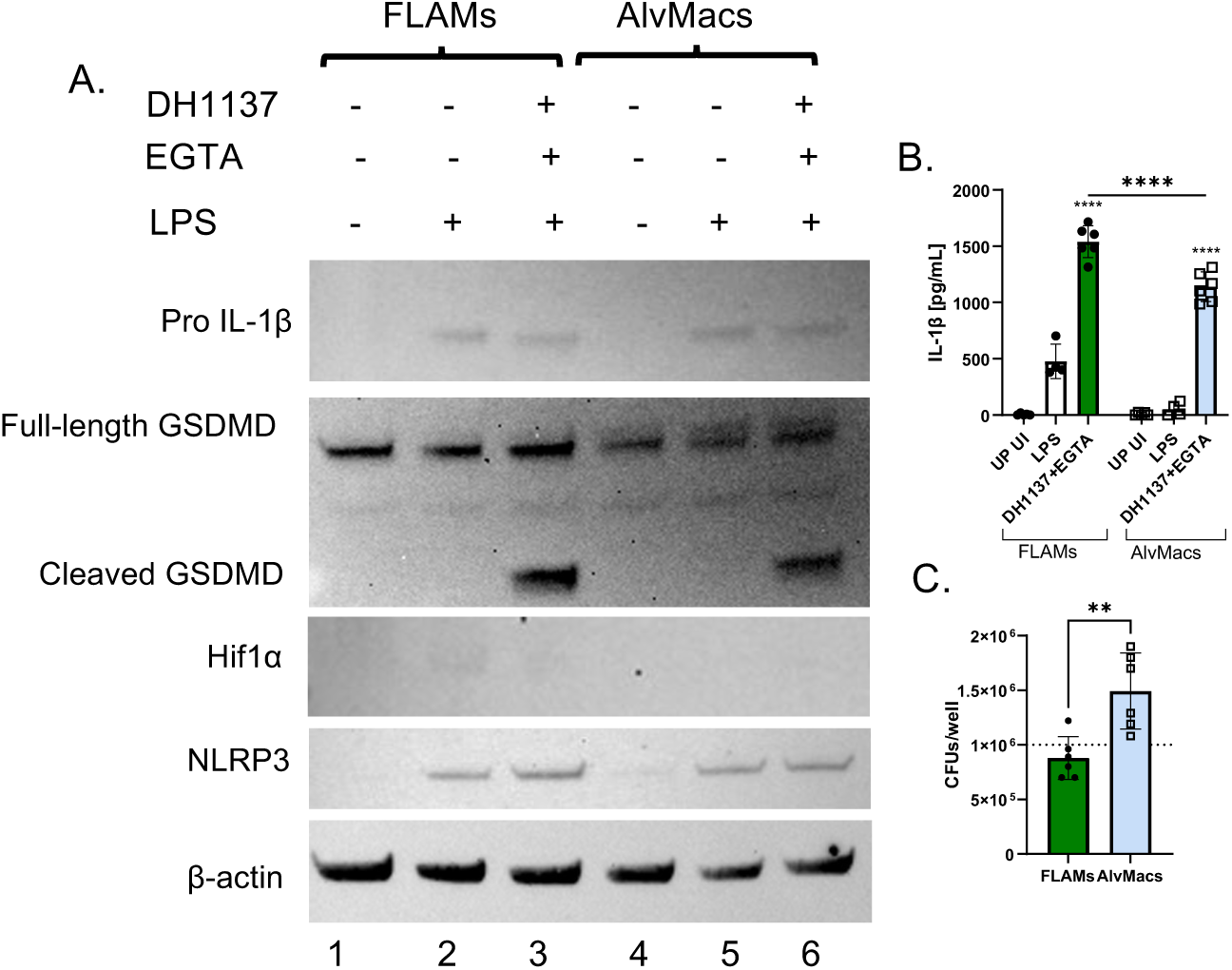
DH1137 infection outcomes in FLAMs or AlvMacs. FLAMs or AlvMacs were left unprimed or primed with LPS for 18 hrs and left uninfected or infected at a MOI=1 for 1.5 hours with DH1137 following subculture in the absence or presence of EGTA. (A) Western blotting of FLAM cell lysates was performed to probe for the indicated proteins. Results are representative of 3 Independent experiments. (B) IL-1β levels in supernatants were quantified via ELISA. (C) Viable bacteria in the combined supernatant and cellular fraction was quantified by CFU assay, with input of 1x106. In (B,C) data is pooled from 3 or more independent experiments where each point represents a single well, and bars show the mean ±SD. (B) Statistical significance was determined using two-way ANOVA with Sidaks multiple comparisons post test comparing to LPS alone or between conditions as shown by brackets. (C) Statistical significance was determined using a student’s *t*-test. (****p<0.0001, **p<0.01).

### Hypoxia Reduces Inflammasome Activation by *P. aeruginosa* in FLAMs and AlvMacs

Given that DH1137 is adapted to hypoxia, we next investigated how a low-oxygen environment influences priming and infection responses in FLAMs. FLAMs were primed with LPS for 5 hrs in normoxia (∼21% O2) or hypoxia (1%) prior to infection with DH1137. Hypoxia during priming reduced production of pro-IL-1β and increased Hif1α (**Figure 5A,B**). Elevated HIF1α confirmed successful cellular adaptation to the oxygen-limiting environment. Because NLRC4 is the primary inflammasome activated in macrophages during *P. aeruginosa* infection (36–38), we probed for total NLRC4 levels but found no differences between the two oxygen environments (**Figure 5B**). Surprisingly, hypoxia reduced cleavage of GSDMD and significantly suppressed IL-1β secretion levels in infected FLAMs (**Figure 5A,C**). LDH release assay showed that cytotoxicity was equivalent in normoxia and hypoxia **(Figure 5D**).

**Fig. 5.**
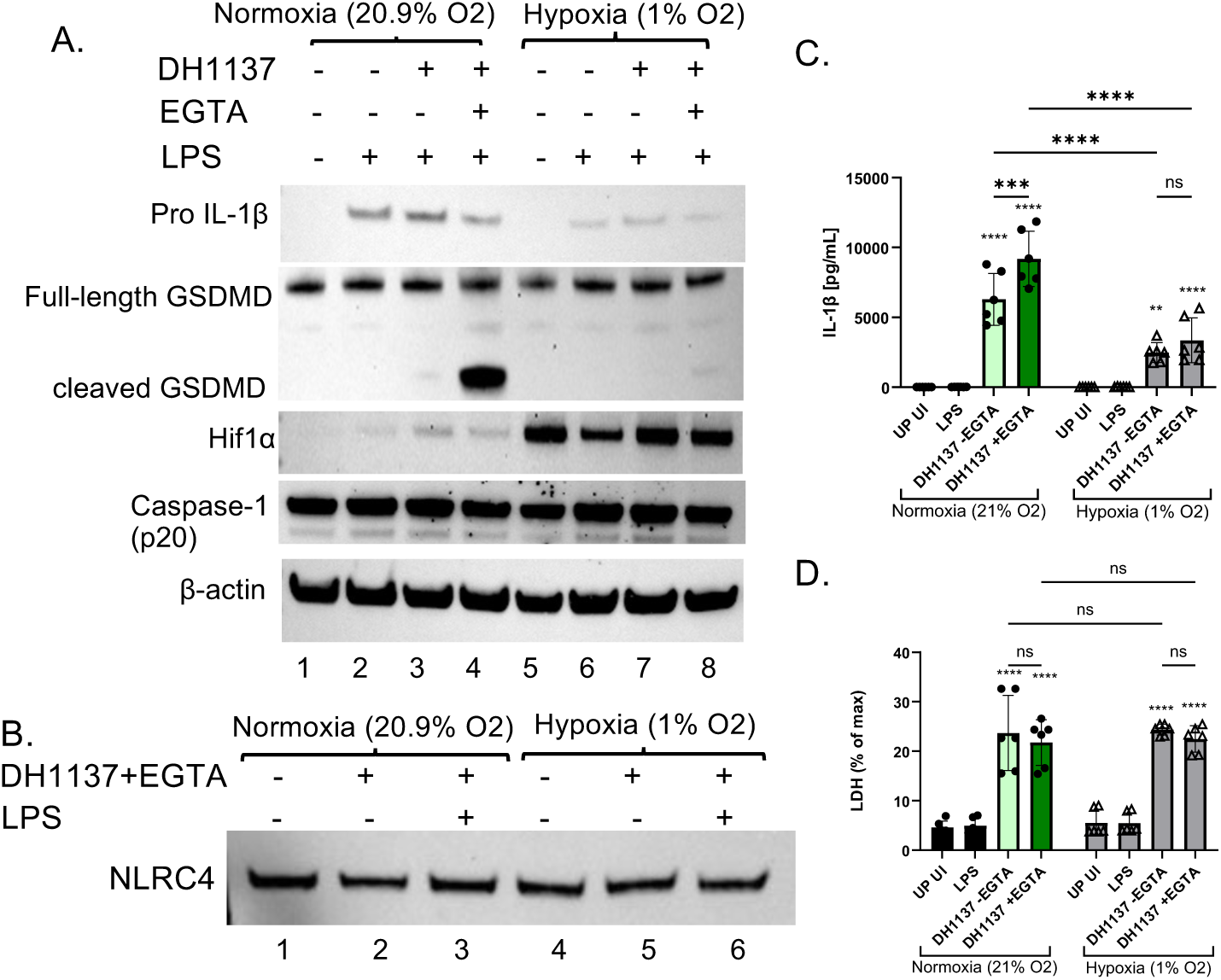
DH1137 infection outcomes in FLAMs in normoxia or hypoxia. FLAMs were left UP or primed with LPS in either in normoxia or hypoxia for 5 hrs and and left UI or infected in the same environment at a MOI=10 for 1.5 hours with DH1137 following subculture in the absence or presence of EGTA. (A,B) Western blotting of FLAM lysates was performed to probe for the indicated proteins. Results are representative of 3 Independent experiments. (C) IL-1β levels in supernatants were quantified ELISA. (D) FLAM pyroptosis was quantified a percentage of maximum LDH release. In (C,D) data is from 3 or more independent experiments where each dot represents a single well, and bars show the mean ±SD. Statistical significance was determined using a two-way ANOVA with Sidaks multiple comparisons post test comparing to LPS alone or between conditions as shown by brackets. (****p<0.0001, ***p<0.001, **p<0.01, *p<0.05, ns not significant).

To determine if hypoxia reduces inflammasome activation by DH1137 in AlvMacs, we compared them to FLAMs. Western blotting confirmed that hypoxia increased production of Hif1α in AlvMacs during priming (**Figure 6A**). In addition, GSDMD cleavage and mature IL-1β release were diminished in hypoxia (**Figure 6A, B**). Interestingly, hypoxic infection with DH1137 yielded significantly higher bacterial loads for both FLAMs and AlvMacs (**Figure 6C**). This finding fits with our previous finding that DH1137 replicates more efficiently in low oxygen (**Figure S2**) and indicates that reduced inflammasome activation is not due to lower bacterial viability in hypoxia. In addition, hypoxia has been shown to impair macrophage bactericidal activity (93). Infection of FLAMs and AlvMacs with PAO1 also resulted in significantly lower IL-1β release from both types of cells in hypoxia (**Figure 6D**). Taken together, these data demonstrate that a low-oxygen environment dampens inflammasome activation in FLAMs and AlvMacs following infection with *P. aeruginosa*, which ultimately correlates with increased survival of DH1137.

**Fig. 6.**
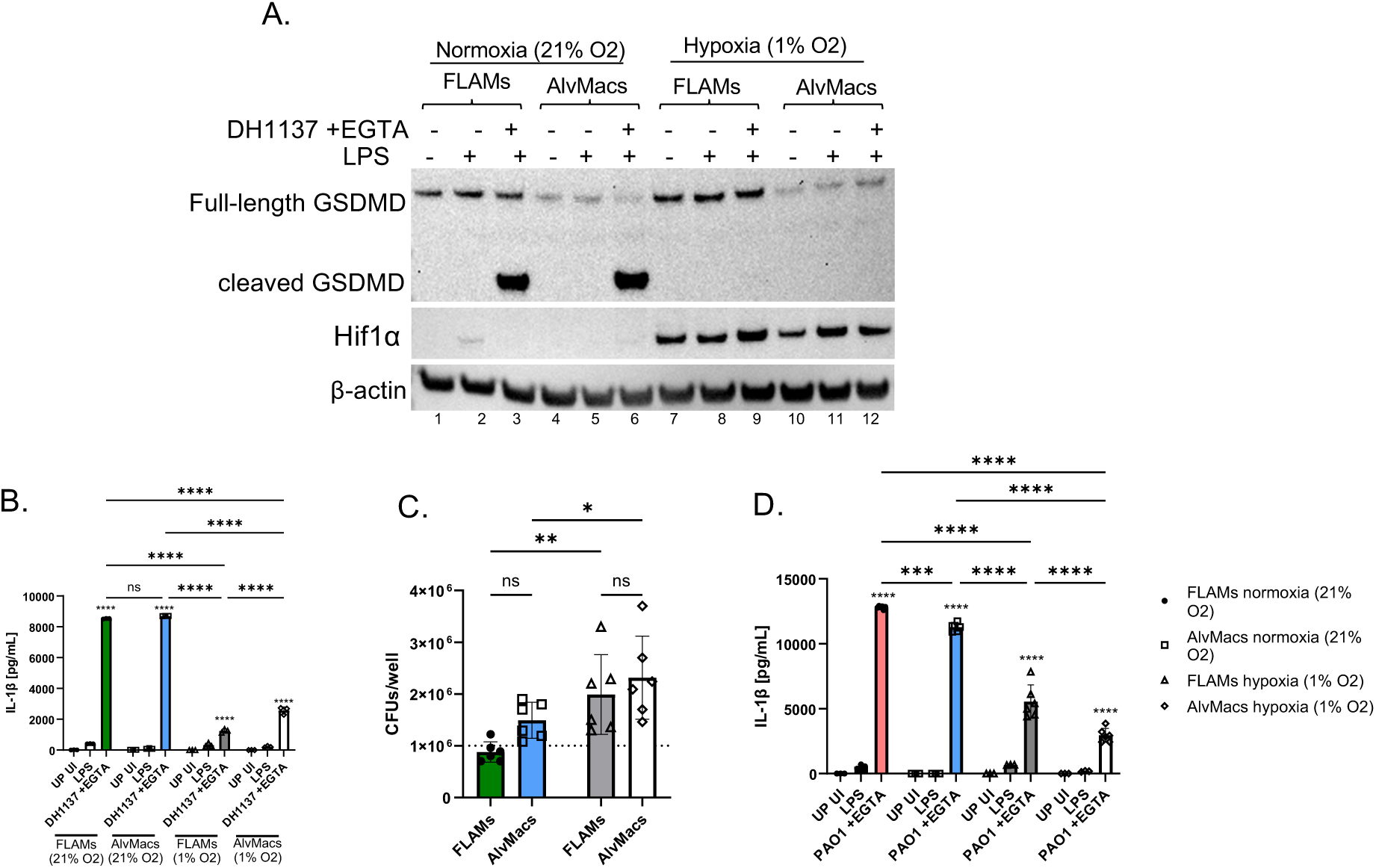
*P. aeruginosa* infection outcomes in FLAMs or AlvMacs in normoxia or hypoxia. FLAMs or AlvMacs were left UP or primed with LPS in either in normoxia (21% O2) or hypoxia (1% O2) for 5 hrs and subsequently infected in the same environments with DH1137 or PAO1 at a MOI=1 for 1.5 hrs following subculture in the presence of EGTA. (A) Western blotting of FLAM cell lysates was performed to probe for the indicated proteins. Results are representative of 3 Independent experiments. (B,D) IL-1β levels in supernatants were quantified via ELISA. (C) Viable bacteria in the combined supernatant and cellular fraction was quantified by CFU assay, with input of input of 1x106. In (B-D) data is pooled from 3 or more experiments where each dot represents a single well, and bars show the mean ±SD. (B,D) Statistical significance was determined using a two-way ANOVA with Sidaks multiple comparisons post test comparing to LPS alone or between conditions as shown by brackets. (C) Statistical significance was determined using a student’s *t*-test. (****p<0.0001, ***p<0.001, **p<0.01, *p<0.05, ns not significant).

### Hypoxia Increases Inflammasome Activation by Nigericin in FLAMs

Having established that hypoxia dampens the inflammasome response to *P .aeruginosa* in alveolar macrophages, we next sought to determine if this outcome extends to other stimuli. FLAMs primed with LPS in normoxia or hypoxia were stimulated with Nigericin—a potent potassium/proton (K^+^/H^+^) antiporter that activates the NLRP3 inflammasome (94). Interestingly, FLAMs treated with Nigericin in hypoxia exhibited elevated levels of cleaved GSDMD compared to those in normoxia (**Figure 7A**). This stood in stark contrast to FLAMs infected with DH1137, which showed drastically reduced GSDMD cleavage under hypoxic conditions. An opposing trend was also observed with mature IL-1β release, with hypoxia causing a significant increase with Nigericin and a reduction with DH1137 (**Figure 7B**). Together, these findings indicate that in alveolar macrophages hypoxia does not inhibit caspase-1 activation, maturation of IL-1β, or GSDMD cleavage per se. Rather, it appears that hypoxia in alveolar macrophages reduces the ability of NAIP/NLRC4 to detect *P. aeruginosa* or assemble active inflammasomes.

**Fig. 7.**
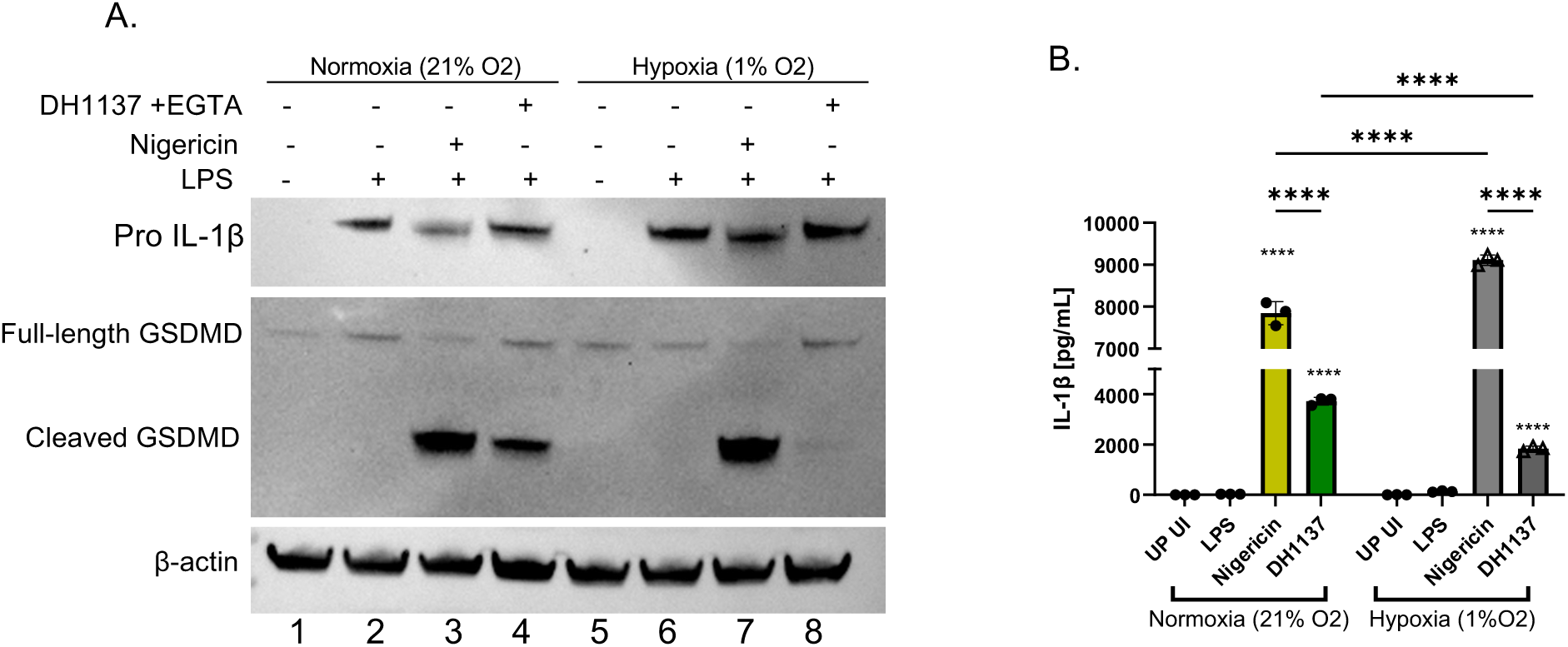
DH1137 infection or nigericin treatment outcomes in FLAMs in normoxia or hypoxia. FLAMs were left UP or primed with LPS for 5 hrs in normoxia or hypoxia followed by incubation in the same environment with 10μM Nigericin or DH1137 (at an MOI=1 following subculture in the presence of EGTA) for 1.5 hr. (A) Western blot data blotting of FLAM cell lysates was performed to probe for the indicated proteins. Results are representative of 1 experiment. (B) IL-1β protein levels in supernatants were quantified by ELISA. Data in B is pooled from 3 independent experiments where each dot represents a single well, and bars show the mean ±SD. Statistical significance was determined using a twoway ANOVA with Sidaks multiple comparisons post test comparing to LPS alone or between conditions as shown by brackets. (****p<0.0001, **p<0.01, *p<0.05, ns not significant).

## Discussion

Several phenotypes observed in DH1137 support the idea that this isolate has undergone adaptation to the selective pressures of the CF airway. Unlike the laboratory strain PAO1, DH1137 demonstrated enhanced growth under hypoxic conditions, consistent with adaptation to the oxygen-depleted mucus environment seen in chronic infection (95). DH1137 is part of the *exoS*^+^ Group A of *P. aeruginosa* strains, the clade most frequently isolated from human infections including pwCF (34, 35). DH1137 has a functional T3SS, although production and secretion of ExoS was lower as compared to PAO1, possibly due to one or more codon changes in regulatory genes. In pwCF it has been estimated that 49% of isolates from newly infected children, 18% of isolates from chronically infected children, and 4% of isolates from chronically infected adults retain T3SS function (44). The original paper that describes DH1137 (called NC- AMT0101-2) indicates the isolate came from a young pwCF (96). Interestingly, intracellular ExoS production decreased under hypoxia while production of PcrV and secretion of both proteins was maintained, suggesting that the regulation of T3SS activity in DH1137 is optimized to minimize immune detection while preserving virulence functions. Together with the genomic changes seen among the regulatory genes for T3SS, these findings suggest that DH1137 has evolved coordinated strategies which promote persistence within the CF lung rather than maximizing acute virulence. Our ExoS production and secretion findings with PAO1 also fit with previously published mass spectrometry data with this strain grown in hypoxia (97). Here, ExoS protein levels were increased along with the regulatory transcription factors ExsA, ExsD, and ExsC (97). In addition, hypoxia was shown to increase expression of T3SS genes in PAO1 (98).

DH1137 also exhibited striking oxygen-dependent motility as seen via a plate-based swimming and swarming assay. Prior publications have indicated that hypoxia decreases virulence factor expression in *P. aeruginosa* (99, 100). In addition, there are suggestions in the literature that hypoxia reduces *P. aeruginosa* flagellar motility (77, 101), although our results may be the first direct demonstration of this phenotype. Under normoxic conditions, both swimming and swarming exceeded those of PAO1, whereas hypoxia dramatically reduced motility in DH1137. If these findings can be reproduced with other clinical isolates, they could suggest that oxygen availability may regulate distinct bacterial behavioral states within the CF airway. Prior studies in the pre-HEMT era have demonstrated that isolates from pwCF that are in poor clinical condition correlates with decreased *P. aeruginosa* motility (79). Whether motility in these strains is altered by oxygen levels remains unexplored. During periods of increased oxygen availability, enhanced motility may facilitate *P. aeruginosa* dissemination throughout the lung, increase encounters with epithelial and immune cells, and contribute to the acute inflammatory episodes characteristic of pulmonary exacerbations. Conversely, within hypoxic mucus plugs, reduced motility combined with enhanced bacterial growth may favor long-term persistence. Interestingly, previous work has found macrophages from pwCF to have decreased expression of TLR5, which may help CF isolates like DH1137 to avoid immune detection despite being highly mobile (102). This model is consistent with the heterogeneous oxygen landscape of the CF lung and provides a potential explanation for how chronic *P. aeruginosa* infections transition between relatively quiescent colonization and episodes of acute disease.

Studies with SGM mice have yielded key insights into important roles of the gut microbiota, including impacts on barrier function, immune training, allergy, inflammatory bowel disease and resistance to intestinal infection (82, 103–108). Our study may be the first to employ SGM mice to study the impact of a Cftr genotype and the gut-lung axis on outcomes of airway infection with *P. aeruginosa.* Prior to infection there was decreased abundance of Bacteroides in the SGM of F508del mice as compared to F508del-Het control animals, reflecting the dysbiosis of these bacterial genera in pwCF (83–85). Under the initial DH1137 infection parameters used, there was no significant difference in weight loss or organ burdens between F508del and F508del-Het control mice, although there was a trend toward greater CFU in the former case. Analysis of cytokines in lungs showed that IL-1α and IL-1β had the highest significant differences when comparing all infected mice to uninfected, suggesting that inflammasomes were being activated by DH1137. Levels of IL-1α and IL-1β were not significantly different between F508del and F508del-Het mice, although G-CSF production was enhanced in the former. G-CSF can be produced by a variety of cells and modifies neutrophil proliferation, maturation and function. G-CSF has been found to be elevated in serum and sputum and to correlate negatively with lung disease in pwCF (109).

An important outcome of this work is the validation of FLAMs as an experimentally tractable model for studying Gram-negative bacterial infection in tissue-resident alveolar macrophages. Although BMDMs have been widely used to investigate inflammasome biology, they differ developmentally and functionally from alveolar macrophages (64). RNAseq results demonstrated that LPS priming increases expression of multiple inflammasome-related genes in FLAMs, and Western blotting confirmed production of pro-IL-1β. Upon infection of FLAMs with DH1137 GSDMD was cleaved and IL-1β was released, and these responses were higher with T3SS induction, indicating activation of the NLRC4/caspase-1 inflammasome. Our comparison of FLAMs with both BMDMs and AlvMacs demonstrated that FLAMs more faithfully reproduce the inflammatory phenotypes of resident alveolar macrophages. Similar levels of HIF1α expression, inflammasome activation, bacterial control, and cytotoxicity between FLAMs ad AlvMacs supports the use of the former as a physiologically relevant model. Given the difficulty of maintaining primary alveolar macrophages (110, 111) FLAMs additionally provide an attractive platform for mechanistic studies of pulmonary host-pathogen interactions.

Our findings not only demonstrate that FLAMs and AlvMacs respond to *P.aeruginosa* infection differently from BMDMs, but also that hypoxia profoundly alters these responses. Most notably, infection of FLAMs or AlvMacs with DH1137 or PAO1 under hypoxic conditions resulted in diminished inflammasome activation as observed via reduced IL-1β secretion and increased bacterial survival despite stabilization of HIF1α. Hypoxia is known to decrease macrophage bactericidal activity (93). However, previous studies using BMDMs have reported that hypoxia potentiates inflammasome activation in response to bacterial infection (57). Surprisingly, we observed the opposite in FLAMs and AlvMacs during infection with DH1137 or PAO1 despite production of NLRC4 being maintained in hypoxia. However, sterile activation of NLRP3 with nigericin resulted in enhanced GSDMD cleavage and IL-1β release under hypoxia, demonstrating that FLAMs remain intrinsically capable of robust inflammasome activation in low oxygen. These findings suggest that hypoxia reduces detection of *P. aeruginosa* or activation of the NLRC4 inflammasome in tissue-resident macrophages. Explanations for this effect of hypoxia include reduced T3SS translocation of NAIP ligands, diminished production of NAIPs, or lowered assembly of NLRC4 inflammasomes in alveolar macrophages. In addition to CF AMs having intrinsically reduced bactericidal activity (112), our findings suggest that the hypoxic microenvironment characteristic of the CF airway fundamentally decreases NLRC4 inflammasome activation in these cells and may help explain the persistence of chronic *P. aeruginosa* infection.

Interestingly, our assays that measured cytotoxicity in FLAMs and BMDMs infected with DH1137 yielded unexpected results. LDH release is commonly interpreted as a marker of plasma membrane disruption from inflammasome-dependent pyroptosis downstream of GSDMD pore formation (113–116). In FLAMs or BMDMs infected in normoxia cytotoxicity was not increased by T3SS induction in DH1137, unlike GSDMD cleavage or IL-1β release. In addition, the diminished GSDMD cleavage and IL-1β release we observed in FLAMs during hypoxic infection with DH1137 occurred despite comparable LDH release between normoxia and hypoxia conditions. These results suggest that in addition to being detected by the NLRC4 inflammasome in FLAMs and BMDMs, DH1137 is causing T3SS-independent cytotoxicity.

Although plasma membrane disruption can release pro-IL-1β, we believe that our ELISA primarily detects the mature form, explaining why the GSDMD cleavage and IL-1β data are correlated. *P. aeruginosa* is known to produce other virulence factors that can cause cytoxicity in host cells including Exotoxin A and pyocyanin (1). Additional experiments are needed to understand mechanisms of cytotoxicity in FLAMs and BMDMs infected with DH1137.

Our findings also have broader implications for understanding persistent inflammation in pwCF receiving highly effective CFTR modulator therapies. Although HEMT has substantially improved clinical outcomes, chronic airway inflammation often persists despite reduced bacterial burden. Recent transcriptomic studies have identified AMs as one of the most abundant and transcriptionally altered phagocyte populations in the CF lung post HEMT, with inflammasome-related pathways significantly altered (61). Our results suggest that the hypoxic airway environment itself may be capable of reshaping AM inflammatory responses independently of CFTR dysfunction and that bacterial adaptation to the hypoxic niche can further modify innate immune signaling.

## Materials and Methods

### P. aeruginosa Strains

*P. aeruginosa* strains used in this study were DH1137 (called NC-AMT0101-2), DH1136 (called NC-AMT0101-1) (96), PAO1 (called PAOIF), and PAO1F *pscD* (117). For secretion, motility and in vitro infection assays strains were grown on Luria-Bertani (LB) agar plates, or in LB high salt broth (11.7g/L NaCl) supplemented with 0.2mM MgCl_2_ and 0.5M CaCl_2_ ±5mM EGTA at 37°C (117). DH1137 for mouse infections was grown on LB agar plates, and in regular LB broth at 37°C.

The genomic sequences of DH1137 and DH1136 can be found at BioProject: PRJNA1454894. The genome accession numbers are: DH1136: JBXUEU000000000 DH1137: JBXUET000000000

### ExoS and PcrV Production and Secretion Assay

Overnight (16-hour) cultures of *P. aeruginosa* were sub-cultured 1:100 in fresh LB high-salt broth supplemented with 10mM MgCl_2_ and 0.5mM CaCl_2_ and ±5mM EGTA and grown at 37°C to mid-log phase at OD_600_= 0.5. 1ml of culture was pelleted, supernatant was mixed with Trichloro Acetic Acid (TCA) (final concentration 10%) and incubated overnight at 4°C with shaking. Bacterial pellets were solubilized using BugBuster to recover intracellular proteins and samples were mixed with 1x sample buffer containing DTT. Proteins precipitated with TCA were pelleted by centrifugation, washed with acetone and resuspended in 1x sample buffer containing DTT. Samples of intracellular and secreted proteins were resolved on SDS PAGE gels and Western blotted for either ExoS or PcrV.

### Swim and Swarm Assays

Swim and swarm assays were performed according to previously published methods (118). Semisolid swarming assay media was formulated with precise agar concentrations—typically ranging from 0.4% to 0.7% w/v—and adjusted to a controlled pH. To strictly control experimental variability, exact parameters for nutrient medium volume, plate thickness, and plate dryness were standardized during pouring and curing. Subcultures of DH1137, PAO1 or PAO1*pscD* were prepared as above. Plates were centrally inoculated on or just below the surface with a fixed volume of the fresh bacterial culture to initiate a localized colony zone before being immediately transferred to an incubation chamber set to 37°C and either normoxia or 1% oxygen. Total surface coverage area and branching morphology were visually assessed.

### Mouse Strains

All mice used in this study were in the C57BL/6 genetic background and housed in a Dartmouth CCMR facility.

Conventional mice: C57BL/6J (stock#664) mice were purchased from Jackson Laboratories and female mice aged between 8-12 weeks of age were used for bone-marrow derived macrophage (BMDM) and alveolar macrophage (AlvMac) isolation. Mice with homozygous *Cftr^F508del^* mutations (*Cftr^em1cwr^* or F508del) on the C57BL/6 background were obtained from Case Western Reserve University’s Cystic Fibrosis Mouse Models Core (81). F508del mice were used to derive germfree animals in the Dartmouth Gnotobiotic Core.

Gnotobiotic mice: Conventional SPF F508del mice were used to derive germfree animals in the Dartmouth Gnotobiotic Core via established Caesarian methods using Swiss Webster foster dams (119). Following establishment of a germfree homozygous F508del breeding colony, we backcrossed mice to C57BL/6 to generate heterozygous animals. We then employed a vertical transmission scheme to engraft littermate F508del homozygous and heterozygous pups with synthetic gut microbiota (SGM) (82, 103). We first inoculated germfree F508del heterozygous male and female breeders with SGM (82, 103) immediately prior to commencement of breeding. Briefly, 14 representative human gut bacterial strains (the SGM) were cultured separately in modified yeast short chain fatty acid (mYCFA) broth in a Whitley anaerobic chamber then gavaged twice on consecutive days into mice. Engraftment was confirmed by collection of fecal pellets, extraction of genomic DNA, and 16S rRNA gene amplicon sequencing (120). Three separately housed trio breeding colonies were established and produced multiple litters over a period of approximately 6 months. Following weaning, equal numbers of both male and female F508del homozygous and heterozygous littermates were maintained under gnotobiotic conditions until 8-12 weeks of age, then transferred to SPF facilities prior to the start of infection assays with *P. aeruginosa* DH1137. Equal numbers of both male and female mice aged between 8-12 weeks of age were used for all infections utilizing SGM mice.

### Ethics Statement

All experiments with mice were carried out in accordance with a protocol that adheres to the Guide for the Care and Use of Laboratory Animals of the National Institutes of Health (NIH) and was reviewed and approved (protocol numbers 00002231 and 00002148) by the Institutional Animal Care and Use Committee at Dartmouth College. The Dartmouth College animal program is registered with the U.S. Department of Agriculture (USDA) through certificate number 12-R- 0001, operates in accordance with Animal Welfare Assurance (NIH/PHS) under assurance number D16-00166 (A3259-01) and is accredited with the Association for Assessment and Accreditation of Laboratory Animal Care International (AAALAC, accreditation number 398).

### Pseudomonas aeruginosa Mouse Infection Model

For acute infections with DH1137, SGM mice were anesthetized by isoflurane inhalation and oropharyngeally challenged with 5x10^5^ CFUs in 100μl PBS. Mice were either sacrificed 48 hours post-challenge for cytokine and CFU analysis or monitored for weight loss and survival for 5 days.

### CFU and Cytokine Analysis of Lung Homogenates from Mice

At 48 hours post-challenge *P. aeruginosa* challenge, mice were euthanized by CO_2_ inhalation. Lung homogenate was collected following full lung dissection, separation into left and right, fixation of right lungs and manual homogenization of left lungs using 2mL of cold PBS. Lung homogenate was used for CFU assay by plating serial dilutions on LB agar or PIA plates, incubation at 37C. Remaining homogenate was clarified by centrifugation at 1,000 × *g* for 5 min and stored at -20°C until analysis. Lung homogenate cytokine concentrations were measured by a 32-plex Luminex^®^ assay according to the manufacturer’s instructions and data were acquired on a Luminex 200 system using xPONENT software.

### Fetal Liver-Derived Alveolar Macrophage (FLAM), BMDM, and AlvMac Isolation and Culture

Fetal liver–derived alveolar macrophage-like cells (FLAMs) were generated as previously described (69). Cells were cultured in complete RPMI 1640 supplemented with 10% fetal bovine serum (FBS) (GE) and 1% penicillin-streptomycin. Medium was further supplemented with 30 ng/ml recombinant mouse GM-CSF (PeproTech), and 15 ng/ml recombinant human TGF-β1 (PeproTech).

Bone marrow-derived macrophages (BMDMs) were cultured from the bone marrow of mice and cultured as described previously (91, 121) plus 1% penicillin-streptomycin. After 7 days of differentiation, the BMDMs were seeded at a density of 5x10^5^ cells/well in 24-well plates in 10/10 media containing Dulbecco’s modified Eagle medium (DMEM)+Glutamax (Gibco) containing 10% FBS (GE), 10% L929 cell-conditioned media, 1 mM sodium pyruvate (Gibco), 10 mM HEPES (Gibco).

AlvMacs from the bronchioalveolar lavage fluid (BALF) of C57BL/6J mice were isolated by gently instilling and aspirating 0.5mL of cold PBS repeatedly using a flexible cannula for a total of 2mLs through a small hole made in the trachea. The recovered BALF was immediately placed into FLAM media containing FBS, GM-CSF, TGF-β1, and penicillin-streptomycin all at the same concentrations used for FLAMs.

All cell types were expanded at 37°C in 5% CO2, and cell culture medium was refreshed every 2– 3 days. When cells reached 70–90% confluency, they were detached by incubation for 10 min in cold PBS containing 10 mM EDTA, followed by gentle scraping.

### RNAseq Analysis of Naive and LPS Primed FLAMs

FLAMs were plated in 6-well plates at 1 × 10^6^ cells/well and left untreated or were treated with LPS (*E. coli* - 055:B5 ultrapure Invivogen Cat no. tlrl-pb5lps) for 18 hours. RNA was then isolated using the Direct-zol RNA Extraction Kit (Zymo Research, Cat no. R2072) according to the manufacturer’s protocol. The Illumina Stranded mRNA Library Prep kit (Illumina, Cat no. 20040534) with IDT for Illumina RNA Unique Dual Index adapters was used for library preparation following the manufacturer’s recommendations but using half-volume reactions. Qubit™ dsDNA HS (ThermoFischer Scientific, Cat no. Q32851) and Agilent 4200 TapeStation HS DNA1000 assays (Agilent, Cat no. 5067-5584) were used to measure quality and quantity of the generated libraries. The libraries were pooled in equimolar amounts, and the Invitrogen Collibri Quantification qPCR kit (Invitrogen, Cat no. A38524100) was used to quantify the pooled library. The pool was loaded onto 2 lanes of a NovaSeq S4 flow cell, and sequencing was performed in a 2×150 bp paired end format using a NovaSeq 6000 v1.5 100-cycle reagent kit (Illumina, Cat no. 20028316). Base calling was performed with Illumina Real Time Analysis (RTA; Version 3.4.4), and the output of RTA was demultiplexed and converted to the FastQ format with Illumina Bcl2fastq (Version 2.20.0).

RNA sequencing (RNA-seq) analysis was completed using the MSU High Performance Computing Center. FastQC (version 0.11.7) was used to assess read quality. Bowtie2 (version 2.4.1) (122) with default settings was used to map reads with the GRCm39 mouse reference genome. Aligned reads counts were assessed using FeatureCounts from the Subread package (version 2.0.0) (123). Differential gene expression analysis was conducted using the EdgeR package (version 4.8.2) in R (version 4.2.1) (124). All raw sequencing data, raw read counts, and normalized read counts will br available through the National Center for Biotechnology Information’s Gene Expression Omnibus database.

### Infection of FLAMs, BMDMs, and AlvMacs with *Pseudomonas aeruginosa*

For *in vitro* stimulation experiments, either 1x10^5^ or 5×10^5^ cells were seeded into 24-well plates and left unprimed or primed with 100 ng/mL O26:B6 *Escherichia coli* LPS (Sigma) and incubated at 37°C with 5% CO_2_. Cells were challenged with live *Pseudomonas aeruginosa* supplied in their respective culture medium mentioned above without 1% penicillin-streptomycin and using phenol-free media (RPMI Corning cat #10-040-CV for FLAMs and AlvMacs and DMEM Gibco cat#10569-010 for BMDMs). Overnight (16-hour) cultures of *P. aeruginosa* were sub-cultured 1:100 in fresh LB high-salt broth supplemented with 10mM MgCl_2_ and 0.5mM CaCl_2_ and ±5mM EGTA and grown at 37°C to mid-log phase at OD_600_= 0.5. Cultures were then pelleted, the LB removed, and bacteria were resuspended in PBS to the original volume. Bacterial suspensions were then diluted to the proper MOI in FLAM infection media. Overnight media from the FLAM cultures was removed and replaced with 100μL of FLAM media containing bacteria. Cell supernatants were collected for cytokine ELISAs and lactate dehydrogenase (LDH) assays. The remaining cells were lysed with M-PER™ (Thermo Scientific) containing DTT (Invitrogen), complete mini (Roche) protease inhibitor, and PhosSTOP (Roche) phosphatase inhibitor. Cell supernatants were removed, and cells were lysed with 100μL 0.1% NP-40 and serially diluted in PBS and plated on LB agar for CFU assays.

For all experiments performed in hypoxia, an InvivO_2_ Hypoxia Chamber (Baker Company/Baker Ruskinn) was used at an oxygen saturation of 1% O2.

### IL-1β Quantification

Murine IL-1β in macrophage culture supernatants was quantified using an ELISA kit (R&D Systems^®^, MLB00C) following the manufacturer’s instructions.

### LDH Quantification

LDH in macrophage culture supernatants was quantified using the CytoTox 96® Non-Radioactive Cytotoxicity Assay (Promega®) following the manufacturer’s instructions.

### Western Blotting

Cell lysates were run on 4-12% NuPAGE Bis-Tris SDS-PAGE gels (Invitrogen by ThermoFisher Scientific) and transferred to PVDF membranes (ThermoFisher Scientific) using an iBlot 2 Gel Transfer Device (Life Technologies). Membranes were blocked in 5% non-fat dairy milk and incubated with primary antibody overnight. The primary antibodies used were rabbit MAb for GSDMD (Abcam, ab209845), rabbit polyclonal for HIF-1 alpha (Novus Biologicals # NB100- 449), mouse monoclonal for caspase-1 (p20) (Adipogen #AG-20B-0042-C100), mouse monoclonal IL-1beta (3A6) (Cell Signaling #12242S), NLRC4 (Cell Signaling #25719T), NLRP3 (Adipogen #AG-20B-0014-C100), rabbit polyclonal for β-actin (Cell Signaling, #4967), rabbit polyclonal antibodies to PcrV and ExoS (from Arne Rietsch). HRP-conjugated anti-rabbit (Jackson Immuno Research) or HRP-conjugated anti-mouse (Jackson Immuno Research) was used as a secondary antibody. Protein bands reacting with antibodies were visualized using chemiluminescent detection reagent (GE Healthcare) on an iBright FL1500 (ThermoFisher Scientific).

## Statistical Analysis of Data

Statistical analyses were performed using GraphPad Prism v11 software, with data representing at least three independent experiments. Group comparisons for ELISA and LDH data were evaluated using either a one-way analysis of variance (ANOVA) with a Tukey post hoc test or a two-way ANOVA with Šídák’s multiple comparisons test. Differences in colony forming units (CFUs) were determined using an unpaired Student’s t-test, while differences in bacterial growth (OD600) between DH1137 and PAO1 under normoxic or hypoxic conditions were analyzed using multiple unpaired t-tests. For non-parametric data, including 32-Plex Luminex® cytokine quantification and in vivo weight loss in SGM mice, significance was determined using the Mann-Whitney test. For RNA-seq analysis, differentially expressed genes were defined using a false discovery rate (FDR) and p-value threshold of <0.05, combined with a log2 fold-change of >1. For all analyses, p-values of ≤ 0.05 were considered significant (*p ≤ 0.01, **p ≤ 0.001, ***p ≤ 0.0001, **** p ≤0.00001, ns: not significant).

## Data Availability

All processed data is included in the manuscript. Raw unprocessed data is being deposited into appropriate databases and accession numbers will be added. The whole genome sequence data and genome assembly are available at BioProject# PRJNA1454894 and genome accession number JBXUET000000000.

## Acknowledgments

We thank members of the Bliska Laboratory for providing feedback on this manuscript. We thank Arne Rietsch for PcrV and ExoS antibodies and Robb Cramer for use of the hypoxia chamber.

---

**Fig. S1.**
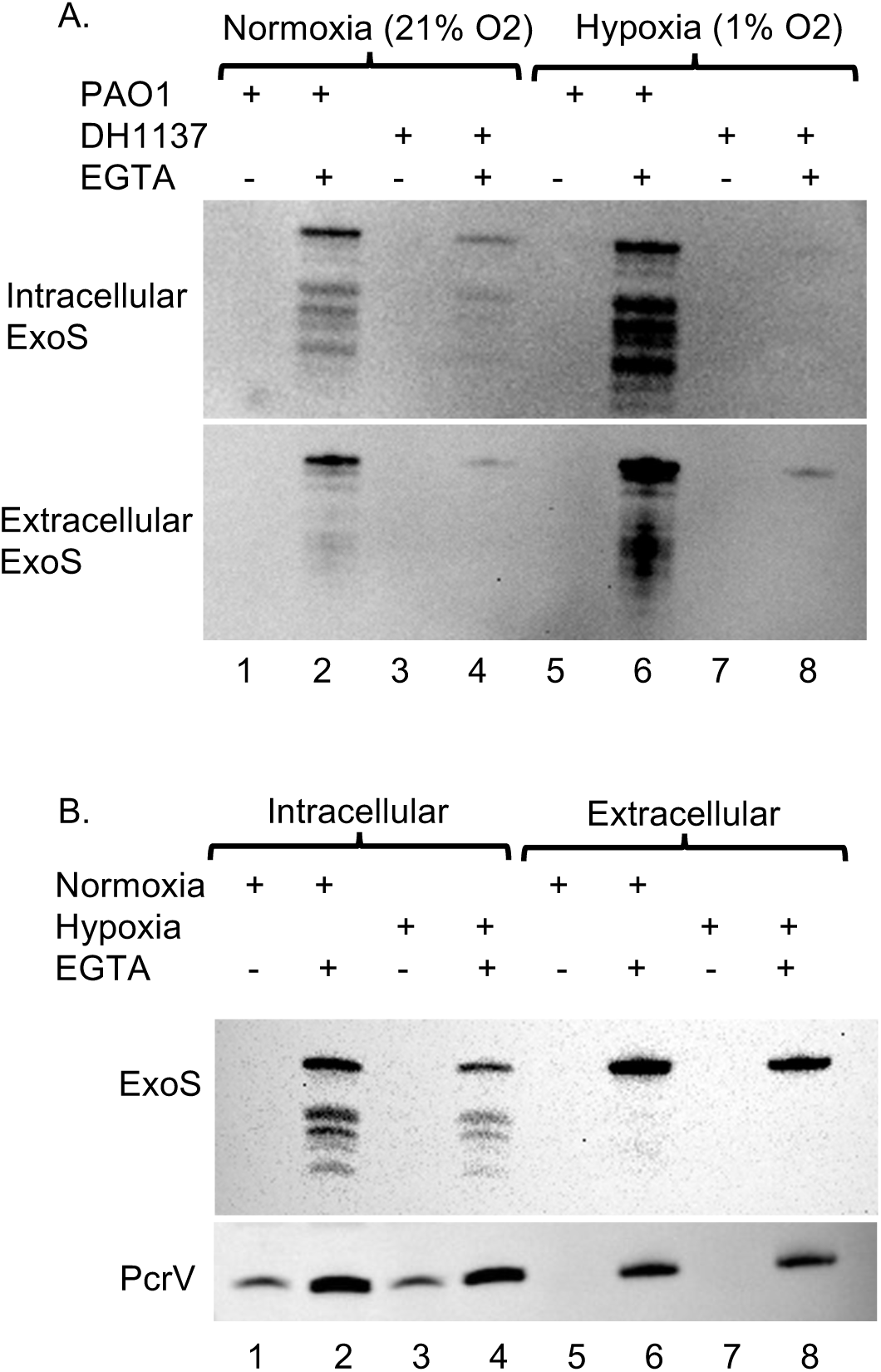
Comparison of ExoS and PcrV production and secretion by PAO1 and DH1137 in normoxia or hypoxia. Normoxia overnights of PAO1 (A) or DH1137 (A,B) were subcultured in LB to the same starting OD600 and grown statically in the absence or presence of EGTA to an OD600=0.5 under normoxia or hypoxia. Western blotting was used to detect intracellular (lysate of cell pellet) or extracellular (precipitate of supernatant) ExoS or PcrV.

**Fig. S2.**
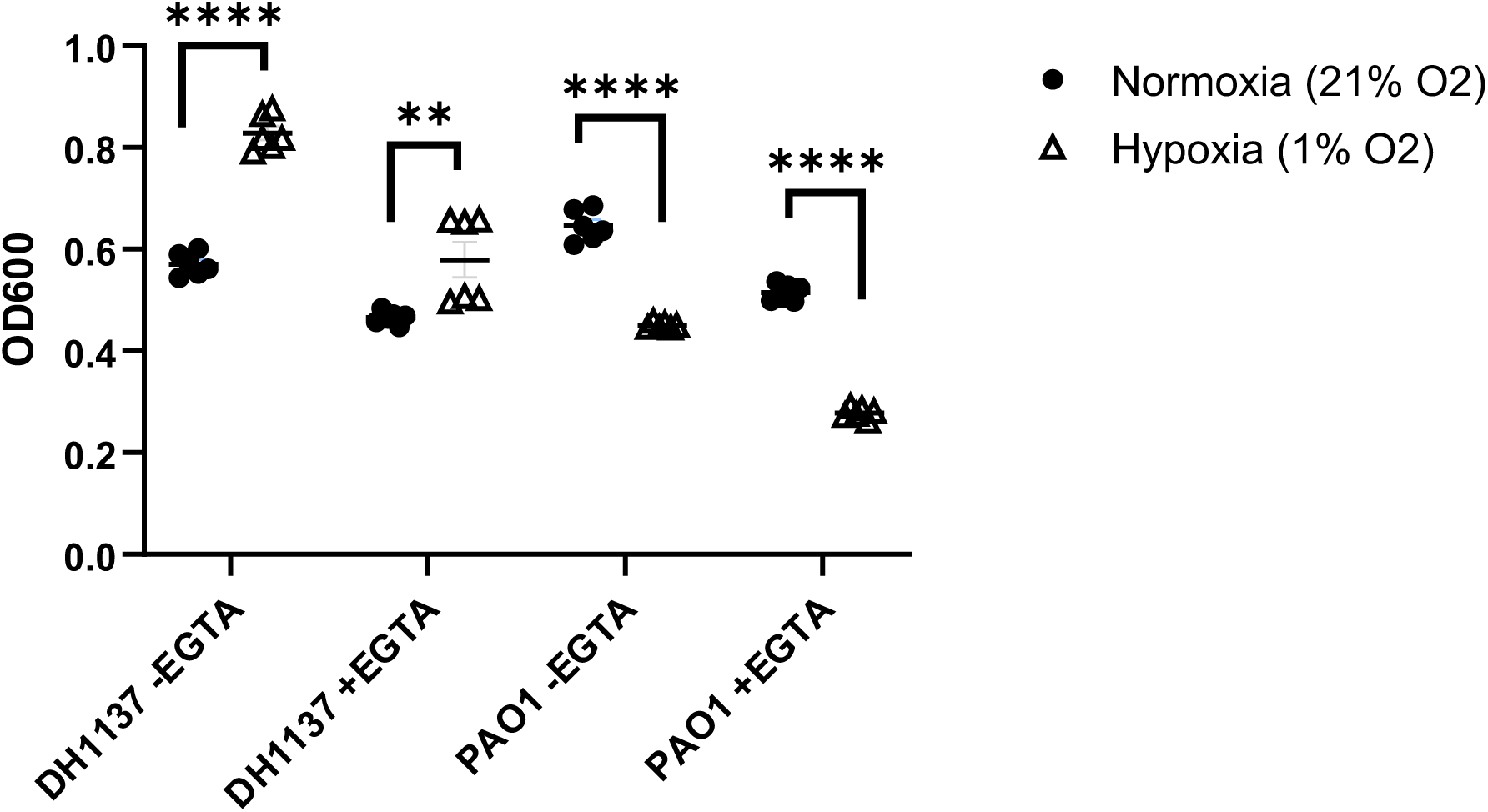
Comparison of DH1137 and PAO1 growth in normoxia or hypoxia. Normoxia overnights of PAO1 or DH1137 were subcultured in LB to the same starting OD600 (approximately 0.05) and grown statically in the absence or presence of EGTA for 4 hrs in normoxia or hypoxia. OD600 values with means were pooled from 2 independent experiments. Statistical significance determined by multiple unpaired t-tests. **p=<0.001; ****p=<0.0001.

**Fig. S3.**
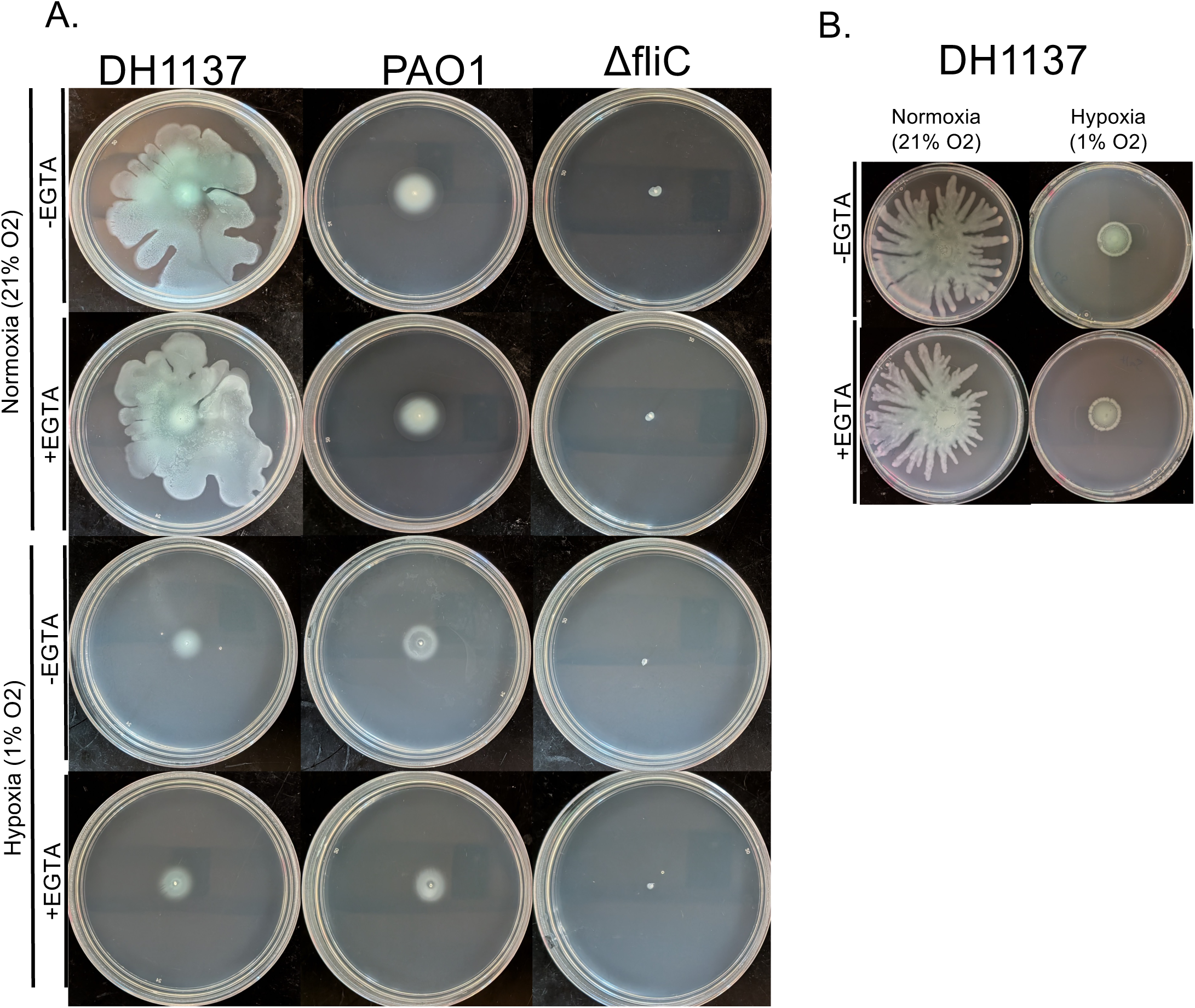
Comparison of DH1137 and PAO1 swim or swarm motility in normoxia or hypoxia. Normoxia overnights of DH1137, PAO1, or PAO1 Δ*fliC* (non-motile control) were subcultured in LB and grown in normoxia in the absence or presence of EGTA and spotted onto swim (A) or swarm (B) agar plates and incubated at 37C in normoxia or hypoxia. Representative images of swim plates after 14 hrs (A) and swarm plates after 18 hrs (B).

**Fig. S4.**
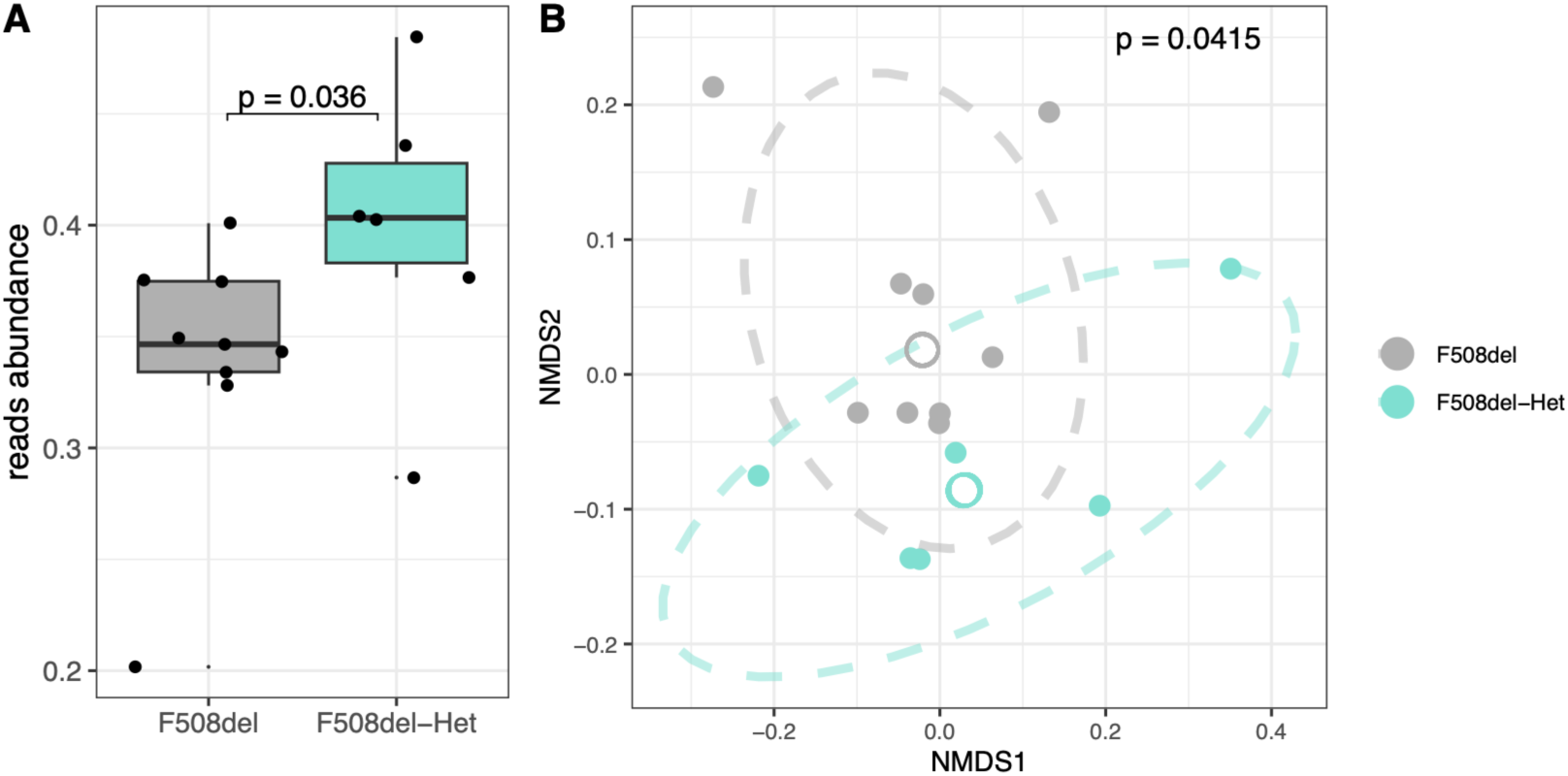
Comparison of vertically transferred SGM in F508del or F508del-Het mice. (A) Comparison of Bacteroides (*B. caccae, B. ovatus, B. thetaiotaomicron,* and *B. uniformis*) abundance between indicated groups (n=9 and n=6, respectively) as determined by V1-V3 16S rRNA gene sequencing and displayed as box and whisker plots. Wilcoxon rank sum test with Bonferroni correction was used to determine p-value. (B) Bray-Curtis NMDS plot comparison of total SGM composition between indicated groups from V1-V3 16S rRNA gene sequencing with Analysis of Similarities (ANOSIM) used to determine p-value. Ellipses indicate 75% confidence intervals. Centroids indicated by hollow points.

**Fig. S5.**
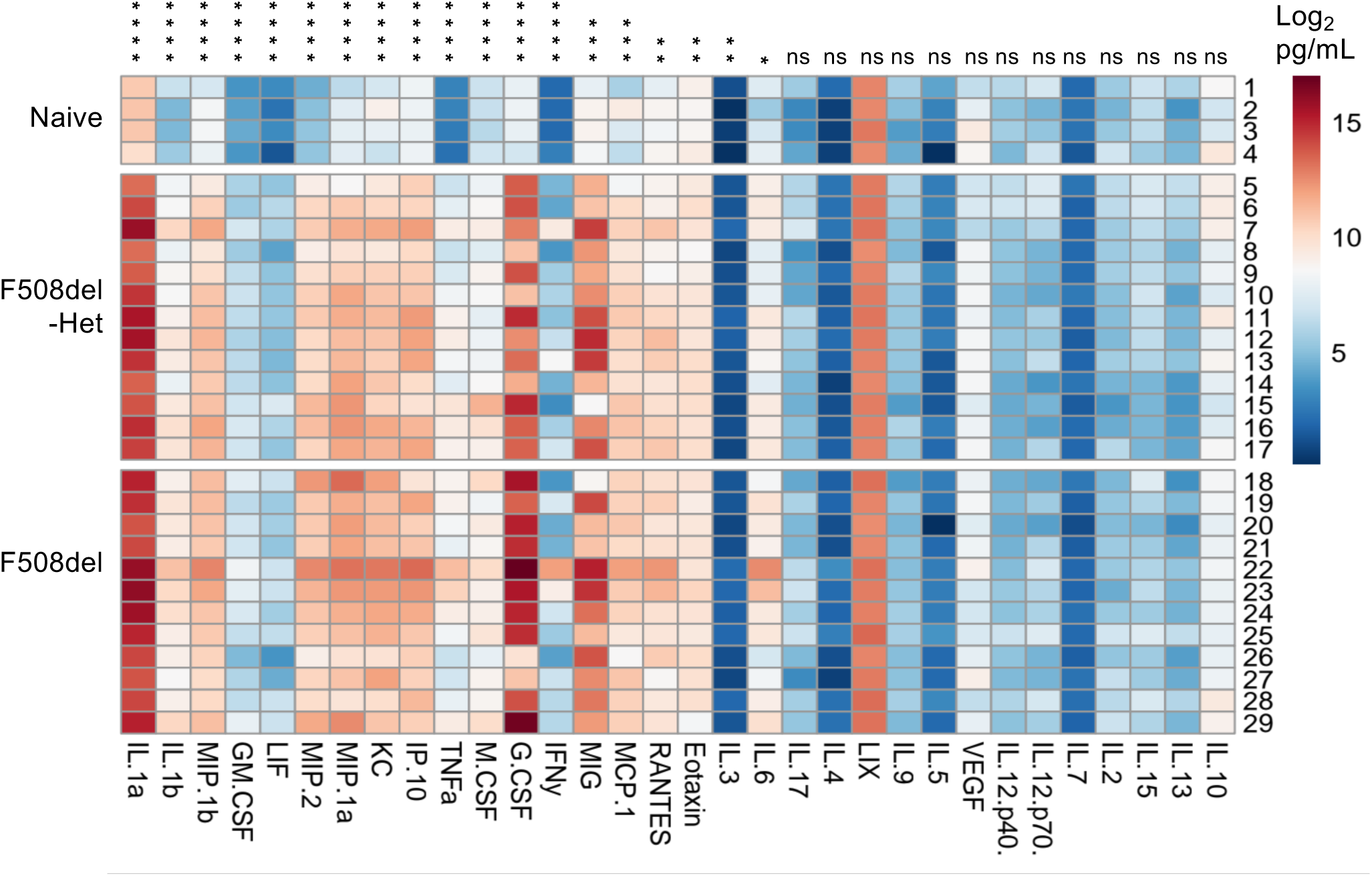
32-Plex Luminex® analysis of lung cytokines in SGM mice. Cytokine levels were measured by 32-Plex in homogenates of right lung lobes from the indicated mice at 48 hrs post mock infection (naïve) or DH1137 infection. Data is presented as heat maps of log2 pg/mL pooled from 3 independent experiments, n=4 for naïve, n=13 for F508del-Het and n=12 for F508del. The genotypes of the naïve mice were F508del-Het (#1,3), wild-type (#2) and F508del (#4). Cytokine order from left to right was determined by statistical significance of naïve compared to all infected mouse samples by a Mann-Whitney test, ****P<0.0001; ***P<0.001; **P<0.01; *P<0.05, ns not significant.

**Fig. S6.**
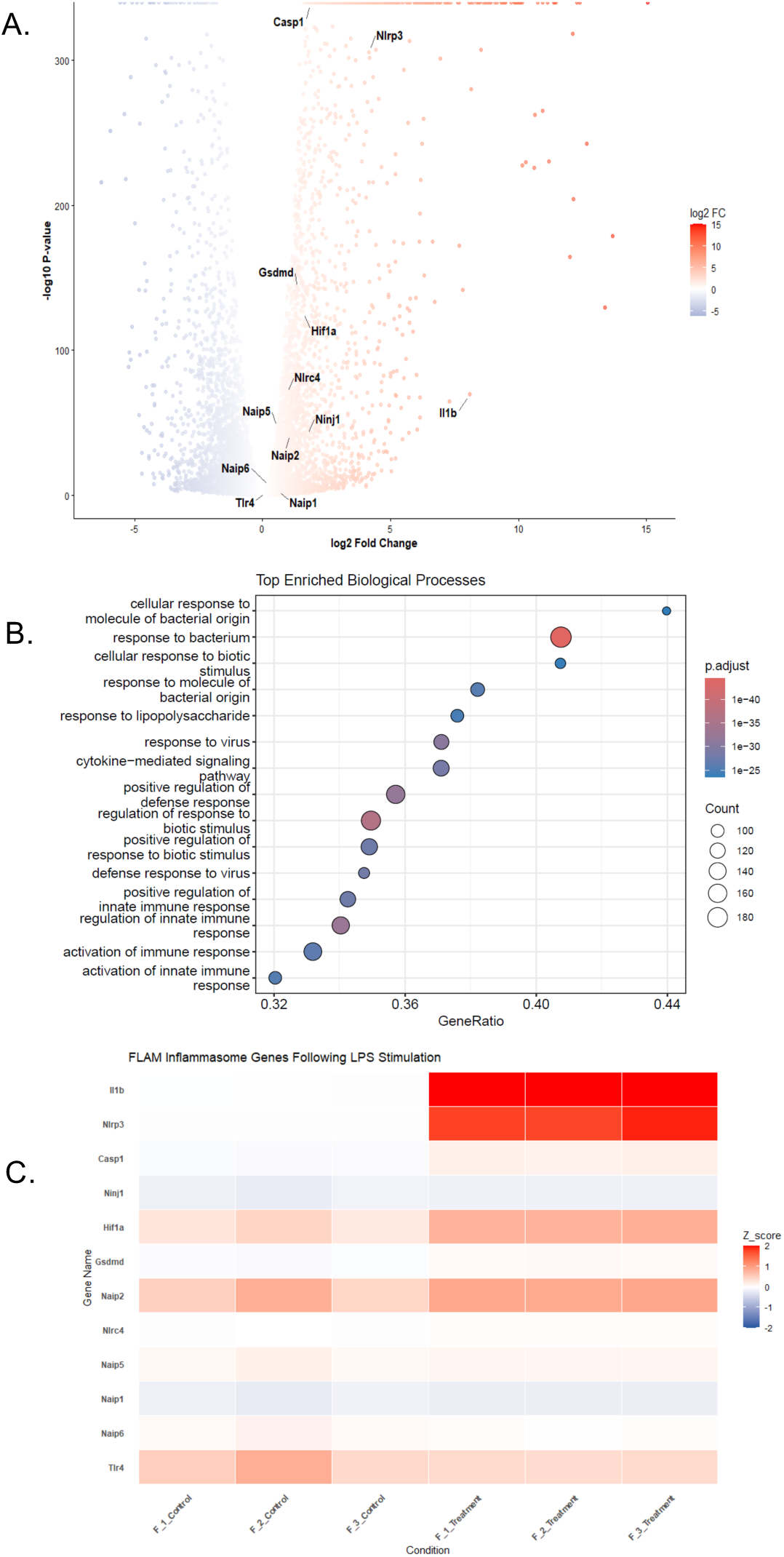
Gene expression analysis of FLAMs primed with LPS. Bulk RNA sequencing analysis was performed on FLAMs incubated with LPS (50ng/mL *E. coli* O111:B4) for 18 hrs. (A) Volcano plot showing which genes are differentially upregulated (red) and differentially downregulated (blue). In summary, 2,046 upregulated and 2,075 downregulated genes were determined with a false-discovery rate and p-value threshold <0.05 and a log2 fold-change >1. Inflammasome-related genes are indicated. Data shows the average of 3 technical replicates from one experiment. (B) Gene Ontology (GO) enrichment analysis showing the top enriched biological processes after incubation with LPS. Adjusted p value (p.adjust) is shown by color and transcript reads (Count) are shown by circle size. (C). Heatmap of inflammasome-related gene expression in FLAMs following an 18 hr LPS stimulation. Ordered from top to bottom on fold change. Relative coloration is standardized by z-score.

**Fig. S7.**
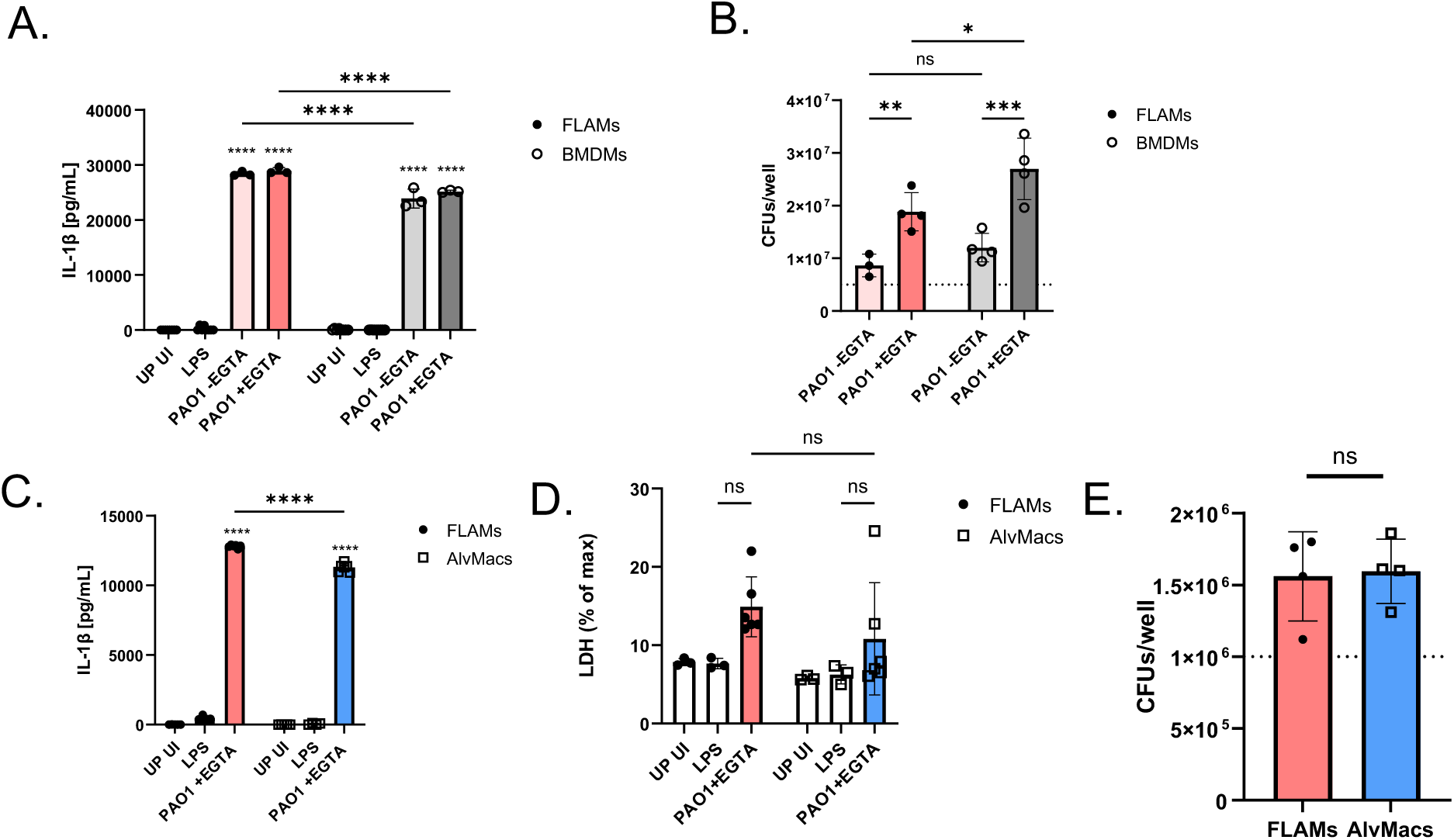
PAO1 infection outcomes in FLAMs, BMDMs or AlvMacs in normoxia or hypoxia. FLAMs, BMDMs or AlvMacs were left UP or primed with LPS for 18 hrs (A-E) or 5 hrs (F) were left UI or infected with PAO1 following subculture in the absence or presence of EGTA at a MOI=10 (A,B) or 1 (C-F) for 1.5 hrs in normoxia (A-E) or normoxia vs hypoxia (F). (A,C,F) IL-1β levels in supernatants were quantified via ELISA. (D) FLAM or AlvMac pyroptosis was quantified as a percentage of maximum LDH release. (B,E) Viable bacteria in the combined supernatant and cellular fraction was quantified by CFU assay, with input of 5x10^6^ (B) or 1x10^6^ (E). Data is pooled from two or more independent experiments where each dot represents a single well, and bars show the mean ±SD. In (A,C,D,F) statistical significance was determined using a two-way ANOVA with Sidaks multiple comparisons post test comparing to LPS alone or between conditions as shown by brackets. In (B,E) statistical significance was determined using a student’s *t*-test.(****p<0.0001, ***p<0.001, **p<0.01, *p<0.05, ns not significant).

**Table S1.** Codon changes in DH1137* vs PAO1

| Gene | Codon change(s) | Protein function | Notes |
| --- | --- | --- | --- |
| PAO763_mucA | T120A | Negative regulator of AlgU, negative regulation of mucoidy and positive regulation of T3SS | Amino acid 120 is in the region where MucA interacts with AlgU. |
| PA1712_exsB | Q105R | Pilotin for T3SS assembly |  |
| PA1714_exsD | A227T, D247E | Negative regulator of ExsA, negative regulation of T3SS |  |
| PA3974_ladS | V509A | Sensor kinase, negative regulation of T3SS |  |
| PA4856_retS | A46V | Sensor kinase, positive regulation of T3SS |  |

\*DH1137 is called NC-AMT0101-2 in Table 1 of Hoffman et al. PMID: 20072604.
https://pmc.ncbi.nlm.nih.gov/articles/PMC2795201/. DH1137 is a CF clinical isolate collected from the same patient as young patient NC-AMT0101-3 but 8.6 years later with wild-type *lasR* allele and mucoid phenotype. The multi locus sequence type (MLST) of DH1137 is ST1209. MLST type 1209 has the following alleles: *acsA*(121), *aroE*(4), *guaA*(26), *mutL*(11), *nuoD*(4), *ppsA*(38), *trpE*(9).

